# CaV1 and CaV2 calcium channels mediate the release of distinct pools of synaptic vesicles

**DOI:** 10.1101/2022.05.03.490438

**Authors:** Brian D. Mueller, Sean A. Merrill, Shigeki Watanabe, Ping Liu, Anish Singh, Pablo Maldonado-Catala, Alex Cherry, Malan Silva, Andres Villu Maricq, Zhao-Wen Wang, Erik M. Jorgensen

## Abstract

Activation of voltage-gated calcium channels at synapses leads to local increases in calcium and the fusion of synaptic vesicles. However, presynaptic output will be determined by the density of calcium channels, the dynamic properties of the channel, the distance to docked vesicles, and the release probability at the docking site. We demonstrate that at *C. elegans* neuromuscular junctions two different classes of voltage-gated calcium channels, CaV2 and CaV1, mediate the release of distinct pools of synaptic vesicles. CaV2 channels are concentrated in densely packed clusters ∼300 nm in diameter with the active zone proteins Neurexin, α-Liprin, SYDE, ELKS/CAST, RIM-BP, α-Catulin, and MAGI1. CaV2 channels mediate the fusion of vesicles docked adjacent to the dense projection and are colocalized with the synaptic vesicle priming protein UNC-13L. By contrast, CaV1 channels are dispersed in the synaptic varicosity and are coupled to internal calcium stores via the ryanodine receptor. CaV1 and ryanodine receptor mediate the fusion of vesicles docked broadly in the synaptic varicosity and are colocalized with the vesicle priming protein UNC-13S. Distinct synaptic vesicle pools, released by different calcium channels, could be used to tune the speed, voltage-dependence, and quantal content of neurotransmitter release.

## Introduction

Synaptic vesicles fuse to the plasma membrane within the presynaptic bouton in a domain called the active zone, and the intricate molecular architecture within the active zone determines the dynamics of the neurotransmitter release (Guzikowski & Kavalali, 2021). Vesicle fusion is driven by calcium influx and binding to the calcium sensor synaptotagmin on the synaptic vesicle (Geppert et al., 1994; Littleton, Stern, Schulze, Perin, & Bellen, 1993; Ward, Weber, & Chapman, 2004). The coupling of fusion sites to calcium channels determines the transfer function of synapses to depolarizing inputs (Eggermann, Bucurenciu, Goswami, & Jonas, 2012; Eguchi, Montanaro, le Monnier, & Shigemoto, 2022; Özçete & Moser, 2021; Rebola et al., 2019), and thus synaptic activity depends on three features of calcium influx: the dynamic properties of the calcium channel, the concentration of calcium at the fusion site, and the release probability of the vesicle. These features are dictated by calcium channel type and location, by the distance to docked vesicles, and by the activity of the priming protein Unc13. Here, we characterize these features at the *C. elegans* neuromuscular junction.

Voltage-gated calcium channels can be divided into three molecular families: CaV1, CaV2, and CaV3, each with fundamentally different dynamic properties, including voltage-sensitive activation and inactivation (Catterall, Perez-Reyes, Snutch, & Striessnig, 2005; Nowycky, Fox, & Tsien, 1985). Each of these channel classes is primarily associated with tissue-specific functions: In muscle, CaV1 (L-type) channels mediate contraction and are coupled to the ryanodine receptor to release internal calcium stores (RyR). In neurons, CaV2 (P/Q, N, and R-type) channels drive synaptic transmission. In neurons and excitable cells, CaV3 (T-type) regulate action potential oscillations and pacemaker frequencies (Dolphin, 2021) These tissue-specific roles are not exclusive, for example, the CaV1 variants CaV1.3 and CaV1.4 are associated with neurotransmitter release in hair cells and photoreceptors, respectively (Dolphin & Lee, 2020).

In the nematode *C. elegans*, each class is encoded by a single gene: CaV1 (*egl-19*), CaV2 (*unc-2*), CaV3 (*cca-1*), and RyR (*unc-68*). In all animals, CaV2 is the main calcium channel for synaptic transmission (Richmond, Weimer, & Jorgensen, 2001; Smith et al., 1996; Tsien, Lipscombe, Madison, Bley, & Fox, 1988; Tsien & Tsien, 1990; Zheng et al., 1995). Unlike other animals, nematodes lack voltage-gated sodium channels and neurotransmission is mediated via graded release (Liu, Chen, & Wang, 2014; Liu, Hollopeter, & Jorgensen, 2009). In *unc-2* mutants, which lack the CaV2 channel, the frequency of tonic miniature currents (’minis’) is severely reduced, but some release remains (Richmond et al., 2001; Tong et al., 2017). Physiological studies suggest CaV1 can also contribute to neurotransmission; CaV1 channel blockers reduce tonic minis (Tong et al., 2017). However, the role of CaV1 channels at synapses in *C. elegans* is complicated because CaV1 also contributes to calcium-mediated action potentials in neurons and is required in the body muscle for viability (Lee, Lobel, Hengartner, Horvitz, & Avery, 1997; Liu, Kidd, Dobosiewicz, & Bargmann, 2018). Finally, the ryanodine receptor also contributes to neurotransmission, and is specifically required for multivesicular release (Chen et al., 2017; Liu et al., 2005).

After the channels close, the concentration of calcium at the site of vesicle fusion will rapidly decline by diffusion (Dittman & Ryan, 2019). Free calcium will be further depleted by calcium buffers and calcium pumps (Blaustein, 1988; Eggermann et al., 2012). Because intracellular calcium is extremely low (0.05 μM), and levels required for fusion are relatively high (half-maximal 10 μM) (Courtney, Briguglio, Bradberry, Greer, & Chapman, 2018; Schneggenburger & Neher, 2000), the effective range of calcium around a single voltage-gated calcium channel is modeled to be only 20 nm for evoked fusion, a ‘nanodomain’ not much larger than the diameter of the calcium channel itself (Fedchyshyn & Wang, 2005; Weber et al., 2010). For synapses to reliably track high frequency action potentials, there must be a large number of channels to negate the stochastic nature of channel opening, and the channels must be tightly coupled to the release sites in this nanodomain. Thus, the transfer function not only depends on the identity of the calcium channel, but also depends on the density of calcium channels and the distance to the docked vesicle. Nevertheless, some synapses rely on ‘loose’ coupling to calcium channels. Calcium microdomains - larger than 100 nm - can drive synaptic vesicle fusion, suggesting that some calcium signals are more robust and do not require tight physical coupling (Eggermann et al., 2012; Vyleta & Jonas, 2014). The tuning of the output of the presynapse then ultimately depends on the spatial and temporal organization of the calcium signal at the release site (Eggermann et al., 2012; Nakamura et al., 2015).

The presence of docked vesicles at release sites and the probability of vesicle fusion depends on the active zone protein Unc13 (Dittman, 2019; Neher & Brose, 2018). Unc13 tethers vesicles to the active zone membrane through C2B and C2C domains which flank the MUN domain (Imig et al., 2014; Quade et al., 2019). The central MUN domain interacts with the SNARE protein syntaxin (Augustin, Rosenmund, Südhof, & Brose, 1999; Lai et al., 2017; Yang et al., 2015) and promotes the open state of syntaxin to initiate SNARE pairing (Richmond et al., 2001). Alternative structural isoforms of Unc13 are distinguished by the presence or absence of a C2A domain at the N-terminus (UNC-13-Long and UNC-13-Short, respectively in *C. elegans*) (Dittman, 2019). Binding of the C2A domain to the scaffolding protein RIM activates these Unc13 isoforms (Betz et al., 2001; Hu, Tong, & Kaplan, 2013; Liu et al., 2019; Lu et al., 2006; Zhou, Stawicki, Goncharov, & Jin, 2013). Unc13 isoforms, which lack a C2A domain, have been demonstrated to bind ELKS / CAST in flies and mice (Böhme et al., 2016; Kawabe et al., 2017). In the absence of Unc13, synaptic vesicles fail to dock at release sites (Hammarlund, Palfreyman, Watanabe, Olsen, & Jorgensen, 2007; Imig et al., 2014; Richmond, Davis, & Jorgensen, 1999; Siksou et al., 2009). Moreover, binding of DAG and calcium to Unc13 regulates the differential release probabilities of primed vesicles (Basu, Betz, Brose, & Rosenmund, 2007; Michelassi, Liu, Hu, & Dittman, 2017; Neher & Brose, 2018).

Here, we demonstrate in *C. elegans* that two different classes of voltage-gated calcium channels, CaV2 (UNC-2) and CaV1 (EGL-19) mediate the release of two physiologically distinct pools of synaptic vesicles. A third calcium channel, the ryanodine receptor (RyR / UNC-68), is essential for CaV1-mediated vesicle release. Time-resolved electron microscopy in mutants demonstrates that these channels mediate fusion of spatially distinct pools of synaptic vesicles in the same synaptic varicosity. Finally, we use super-resolution fluorescence microscopy to demonstrate that CaV2 is localized with UNC-13L at the dense projection, and that CaV1 and RyR colocalize with UNC-13S at distal sites. Altogether, we describe two pools of synaptic vesicles: (1) The central pool is localized adjacent to the dense projection, vesicles are docked by UNC-13L, and released by a dense cluster of CaV2 channels. (2) The lateral pool of vesicles is broadly distributed, docked by UNC-13S, and released by dispersed CaV1 and RyR channels.

## Results

### CaV1 and CaV2 calcium channels are required semi-redundantly for nervous system function

The genome of *C. elegans* contains only a single gene for each major voltage-gated calcium channel class: CaV1 (*egl-19*), CaV2 (*unc-2*), CaV3 (*cca-1*), and a single calcium-gated RyR (*unc-68*) (hereafter, referred to by their common names). Loss of the CaV3 T-type channel does not affect neurotransmitter release in acetylcholine neurons (Liu et al., 2018). However, loss of any other calcium channel results in impaired neurotransmission (Liu et al., 2005; Richmond et al., 2001; Tong et al., 2017). Null mutants lacking either CaV2 (*unc-2*(*lj1)*) or RyR (*unc-68*(*e540*)) are viable, but CaV1 null mutants (*egl-19*(*st556)*) die as embryos due to a loss of muscle function during morphogenesis (Lee et al., 1997). We rescued the CaV1 null mutant using a muscle promoter expressed early in development; since this strain lacks CaV1 in the nervous system, we refer to it as ‘CaV1(Δns)’.

To determine whether these channels function cooperatively or in parallel, we tested for synthetic interactions between mutations of these channel types. The double mutant CaV1(Δns) RyR is viable, and is no worse than the RyR null, consistent with their coupled function (Figure 1A). However, CaV1(Δns) CaV2(−) double mutants and RyR(−) CaV2(−) double mutants exhibit a synthetic lethal interaction. These data suggest that calcium influx from CaV1-RyR acts redundantly, and in parallel, with CaV2 to sustain neurotransmission.

**Figure 1.**
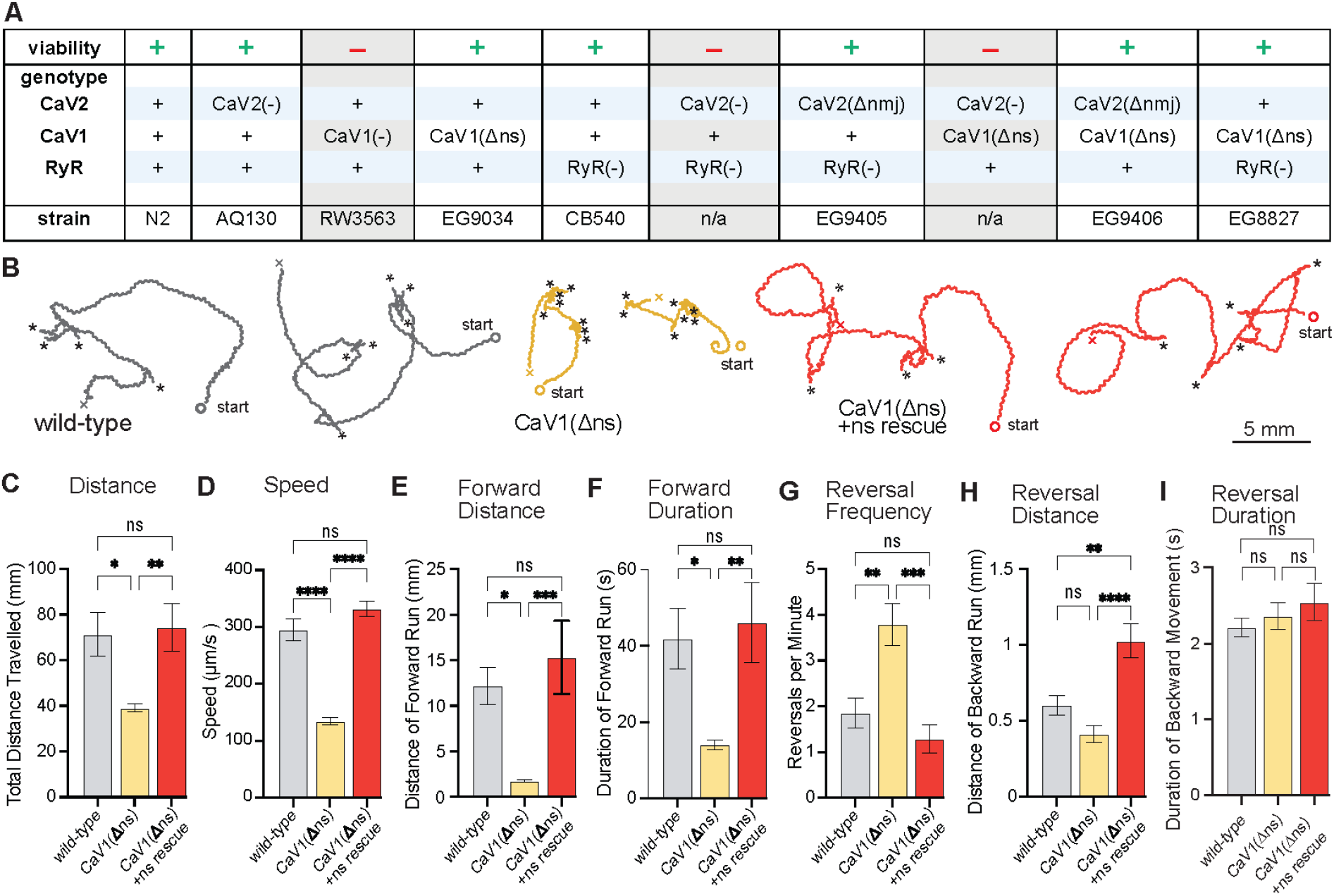
nervous system CaV1 is required for normal behavior and locomotion. (A) Viability of calcium channel double mutants. (B) Worm Tracks. Healthy animals were tracked for 5 minutes with a frame rate of 8 frames per second. The path the animal created was plotted, starting point is indicated, asterisks represent reversal events. (C) Total average distance animals travelled per animal during the 5-minute interval by genotype. Wild-type 71.2 ±9.5 mm. CaV1(Δns) 39.0 ±1.8 mm. CaV1(Δns) +rescue 74.4 ±10.5 mm. (D) Average speed, including both forward and backward bouts but excluding pauses, for the duration of the assay. Wild-type 294.9 um/s ± 19.4 um/s. CaV1(Δns) 133.8 um/s ± 6.3 um/s. CaV1(Δns) +rescue 331.7 um/s ± 13.3 um/s. (E) Average distance of forward locomotion between reversal events that animals travelled by genotype. Wild-type 12.2 mm ±2.0 mm. CaV1(Δns) 17.7 mm ±1.7 mm. CaV1(Δns) +rescue 15.3 mm ± 4.0 mm. (F) Average duration of forward run between reversal events. Wild-type 41.9 ± 8 s. CaV1(Δns) 14.1 ± 1.3 s. CaV1(Δns) +rescue 46.2 ± 10.5 s. (G) Average number of reversal events per minute exhibited by animals by genotype. Wild-type 1.9 ± 0.3 events. CaV1(Δns) 3.8 ± 0.5 events. CaV1(Δns) +rescue 1.3 ± 0.3 events. (H) Average distance travelled in reverse per animal by genotype. Wild-type 601.9 ± 65.1 um. CaV1(Δns) 413.2 ± 56.8 um. CaV1(Δns) +rescue 1026 ± 111.1 um. (I) Average duration of reversal run. Wild-type 2.2 +- 0.1 s. CaV1(Δns) 2.4 +/- 0.2 s. CaV1(Δns) +rescue 2.5 +- 0.2 s. Wild-type n=11, CaV1(Δns) n=16, CaV1(Δns) +rescue n=13. Error bars reported in SEM. Genotypes were blinded. One-way ANOVA with Tukey’s multiple comparisons was used to calculate p-value. *p<0.05, **p<0.005, ***p<0.001, ****p<0.0005 Data available as Source Data 1

CaV1(Δns) animals are uncoordinated, and the phenotypes are fully rescued by the expression of CaV1 in the nervous system (Figure 1B). To determine the role of CaV1 on locomotion, we characterized crawling using worm tracker software. CaV1(Δns) worms travel shorter distances than wild-type worms, consistent with slower crawling speeds (Figure 1C,D). In addition, wild-type animals spent longer time and travelled longer distances during forward bouts (Figure 1E,F). Conversely, CaV1(Δns) worms traveled shorter distances and initiated reversals more frequently than the wild type (Figure 1G). However, the distance travelled while backing tended to be shorter in the CaV1(Δns) animals; whereas the rescued animals travelled longer distances in reverse, about a full body length (Figure 1H). Reversals in all three genotypes were similar in duration (Figure 1I). These results indicate that CaV1 functions in the nervous system for speed of locomotion, and also biases the bistable locomotory circuit toward forward locomotion (Zheng, Brockie, Mellem, Madsen, & Maricq, 1999).

### CaV2 and CaV1 regulate distinct pools of synaptic vesicles

To determine if CaV1 is playing a direct role in synaptic transmission, as well as the depolarization state of the locomotory circuit, we recorded synaptic currents at neuromuscular junctions. Body muscles were voltage-clamped at a holding potential of -60mV, and miniature postsynaptic currents recorded. Miniature postsynaptic currents (‘minis’) are caused by the release of neurotransmitter from one or a few synaptic vesicles. The nematode neuromuscular junction releases neurotransmitter via graded release (Liu et al., 2014; Liu et al., 2009); the frequency of tonic minis drive calcium action potentials in the muscles (Liu et al., 2011).

The rate of miniature postsynaptic currents (minis/s) compared to the wild type (32.2 +/- 2.1 minis/s) is significantly reduced in each of the single mutants, CaV2 (18.7 +/- 3.5 minis/s), CaV1(Δns) (18.7 +/- 3.6 minis/s), and RyR (20.9 +/- 1.5 minis/s) (Figure 2A). Because some of the channel double mutants are synthetic lethal, we acutely blocked CaV1 using nemadipine (Kwok et al., 2006). The frequency of minis in the wild-type with nemadipine (+nema 18.4 +/- 2 minis/s) was similar to CaV1(Δns) and did not further reduce mini frequency in the CaV1(Δns) mutant (+nema 19.1 +/- 2.7 mini/s), demonstrating that nemadipine is an effective blocker of CaV1 and does not block CaV2 nonspecifically. Nemadipine application on the RyR mutant did not exacerbate the phenotype (‘RyR(−) +nema’, 20.7 +/- 1.3 mini/s), indicating that RyR and CaV1 function interdependently at neuromuscular junctions. To determine if CaV1 and CaV2 are required together for all neurotransmitter release, we blocked CaV1 in the CaV2 mutant. Application of nemadipine almost completely abolished mini frequency in the CaV2 mutant (+nema 1.7 +/- 0.6 mini/s). We conclude that all vesicle fusion at neuromuscular junctions relies on CaV1 and CaV2, each contributing about half of the minis. In addition, calcium influx through CaV1 is not sufficient for vesicle fusion; CaV1 relies on internal calcium stores released by RyR to fuse synaptic vesicles.

**Figure 2.**
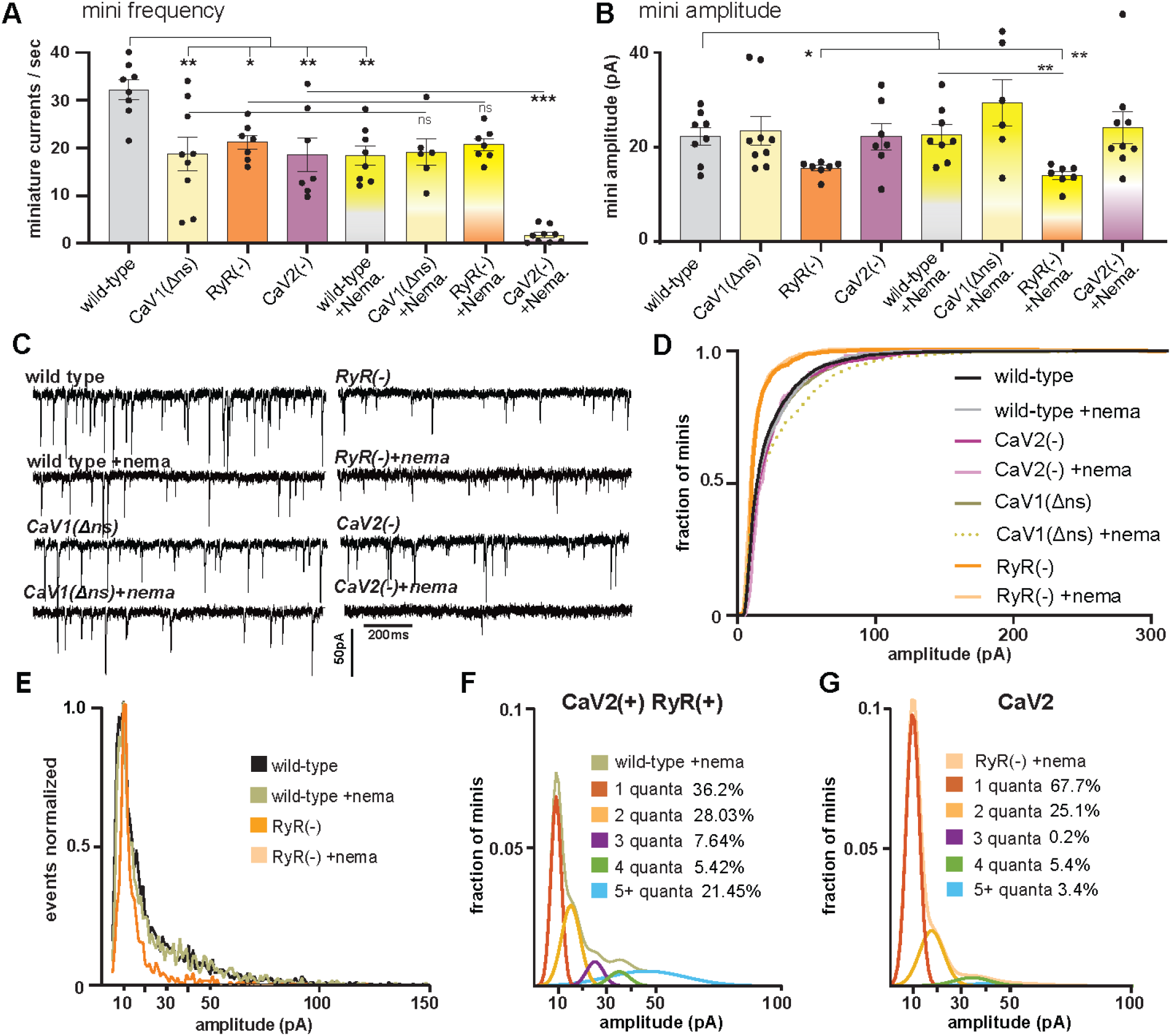
CaV2 and CaV1-RyR fuse different vesicle pools. (A) Spontaneous miniature currents mediated by CaV1 and RyR are inhibited by nemadipine. Wild-type: 32.2 ± 2.1 minis/s n=8, wild type with nemadipine (10 µM): 18.4 ± 2.1 mini/s n=8. CaV1(Δns): 18.7 ± 3.1 minis/s n=9, with nemadipine 19.1 ±2.7 mini/s n=6. RyR(−): 20.9 ± 1.5 minis/s n=7, with nemadipine 20.7 ±1.3 mini/s n=7. CaV2(−): 18.7 ± 3.5 minis/s n=7, with nemadipine 1.7 ± 0.6 mini/s n=9. One-way ANOVA with Dunnett’s multiple comparisons test and one-way ANOVA with Tukey’s multiple comparisons tests were used to calculate significance. (B) The ryanodine receptor mediates large amplitude tonic miniature currents. Wild type 22.3 ± 1.9 pA n=8, wild type with nemadipine 22.6 ± 2.1 pA n=8. CaV2(−): 22.1 ± 3.5 pA n=7, CaV2(−) with nemadipine 24.1 ± 3.4 pA n=9. CaV1(Δns): 23.4 ± 3.1 pA n=9, CaV1(Δns) with nemadipine 29.4 ± .9 pA n=6. RyR(−): 15.5 ± 0.6 pA n=7, RyR(−) with nemadipine 14.0 ± 0.9 pA n=7. One-way ANOVA with Dunnett’s multiple comparisons test and Welch’s t-test were used to calculate significance. (C) Sample traces of spontaneous release in 0.5mM extracellular calcium. (D) Cumulative distribution plot of miniature current amplitudes. (E) Frequency distribution of mini amplitudes of wild-type and ryanodine mutants ± nemadipine, normalized to modal value. (F) Quantal analysis of post-synaptic amplitudes from wild-type ± nemadipine. The cumulative plot of mini amplitudes was replotted into 1 pA bins and a single quantal mini amplitude distribution was fit with a gaussian at a modal value of 9 ±2 pA. The wild-type distribution of amplitudes was fit with a 5-term convolution of a single quanta (khaki-colored curve). 1-quanta (rust) accounted for 36.2% of fusions. 2-quanta (butterscotch) 28.03%. 3-quanta (violet) 7.64%. 4-quanta (green) 5.42%. 5 or more quanta (blue) 21.45%. (G) Quantal analysis of post-synaptic amplitudes from RyR(−) ± nemadipine animals. A 5-term gaussian model was fit to the cumulative frequency of amplitudes 1pA bin size (peach). 1-quanta (rust) accounted for 67.7% of fusions. 2-quanta (butterscotch) 25.1%. 3-quanta (violet) 0.2%. 4-quanta (green) 5.4%. 5 or more quanta (blue) 3.4%. For all recordings, Vm = -60 mV, 0.5 mM calcium. Error bars reported in SEM. *p<0.05, **p<0.005, ***p<0.001, ****p<0.0005. Data available as Source Data 2

We confirmed these conclusions by making viable double mutants. Similar to the nemadipine experiments, the rate of minis in the CaV1(Δns) RyR double mutant (19.6 +/- 2 mini/s) is similar to the single CaV1(Δns) and RyR mutants, again suggesting that CaV1 relies on coupling to RyR to activate neurotransmitter release. To generate strains that bypass the synthetic lethality with CaV2, we expressed CaV2 in head neurons using a tissue-specific promoter (‘P*unc-17*h’) (Hammarlund et al., 2007; Topalidou et al., 2016). Expression of CaV2 in the head neurons of CaV2 nulls, referred to as ‘CaV2(Δnmj)’, bypassed the synthetic lethality of both the CaV1(Δns) CaV2(−) double mutant, and the RyR(−) CaV2(−) double mutant, but the rescued animals exhibit a synthetic paralyzed phenotype. The mini rates of CaV2(Δnmj) CaV1(Δns) double mutants (12.8 +/- 2.4 mini/s) and CaV2(Δnmj) RyR double mutants (11.3 +/- 2.1 mini/s) are significantly diminished, but not completely abolished (Extended Data Figure 2A). The lingering neuronal activity in these strains is likely due to CaV2 expression from head-rescued CaV2 neurons synapsing onto the muscles (the sublateral cord motor neurons). Nevertheless, they demonstrate that CaV1 and RyR are in the same pathway, and act in parallel to CaV2.

### RyR is required for multiquantal release

The mode of mini amplitudes represents miniature currents from single vesicle fusions, and the modal value is similar in all strains (WT 10 pA, CaV1Δns 10pA, RyR 11pA, CaV2 8pA) regardless of nemadipine treatment (N2+nema 10pA, CaV1Δns+nema 8pA, RyR+nema 10pA, CaV2+nema 11pA) (Figure 2E, Extended Data Figure 2D). However, the mean amplitude of miniature currents is reduced in the RyR(−) mutant (15.5 ±0.6 pA) compared to the wild type (22.3 ±1.9 pA; Figure 2B-D, Extended Data Figure 2B,E). The reduction in mini amplitudes is likely due to a loss of multiquantal release since the focus of ryanodine receptor function is in neurons (Chen et al., 2017; Liu et al., 2005).

CaV2 is also functionally coupled to RyR to drive multiquantal release. Mini amplitudes from CaV2 and RyR can be observed when CaV1 channels are blocked by nemadipine (22.6 ±2.1 pA, ‘wild-type + nemadipine’, Fig. 2E). Mini amplitudes solely from CaV2 are reduced in the absence of RyR (14 +/- 0.9pA, ‘RyR + nemadipine’), suggesting that RyR also responds to calcium from CaV2. Further, fitting the frequency distribution of mini amplitudes to multiple gaussians indicate that most multiquantal events are driven by RyR (Figure 2F), though some multiquantal events may be attributed to CaV2 (Figure 2G).

Together, these data demonstrate that CaV2 and CaV1 channels regulate the release of separate synaptic vesicle pools at neuromuscular junctions. CaV1 requires the ryanodine receptor for vesicle fusion. Calcium influx through CaV2 is sufficient to fuse vesicles although neurotransmitter release is amplified by the ryanodine receptor.

### CaV2 and CaV1 mediate fusion of separate vesicle pools at single synapses

The physiology data suggest that CaV2 and CaV1 mediate the release of distinct synaptic vesicle pools onto muscles. To demonstrate these calcium channels regulate spatially distinct pools at the same synaptic varicosity, time-resolved “flash-and-freeze” electron microscopy was used to characterize fusing vesicle pools (Watanabe et al., 2013). Transgenic animals expressing channelrhodopsin in acetylcholine neurons were loaded into a high-pressure freezing chamber, and stimulated with a 20-ms light pulse to depolarize neurons and activate synaptic calcium channels. Animals were frozen 50 ms after stimulation; control animals were treated identically but not stimulated. Frozen samples were fixed by freeze substitution, embedded in plastic and sectioned for electron microscopy (Watanabe et al., 2013). Docked vesicles are defined as those in contact with the plasma membrane; docking was segmented blind to treatment and genotype (Figure 3A,B). The distance from the dense projection to the docked vesicle is plotted on the X-axis (Figure 3C). Decreases in docked vesicles after stimulation are assumed to be the result of synaptic vesicle fusion, although calcium influx could cause some vesicles to undock and return to the cytoplasm (Kusick et al., 2020).

**Figure 3.**
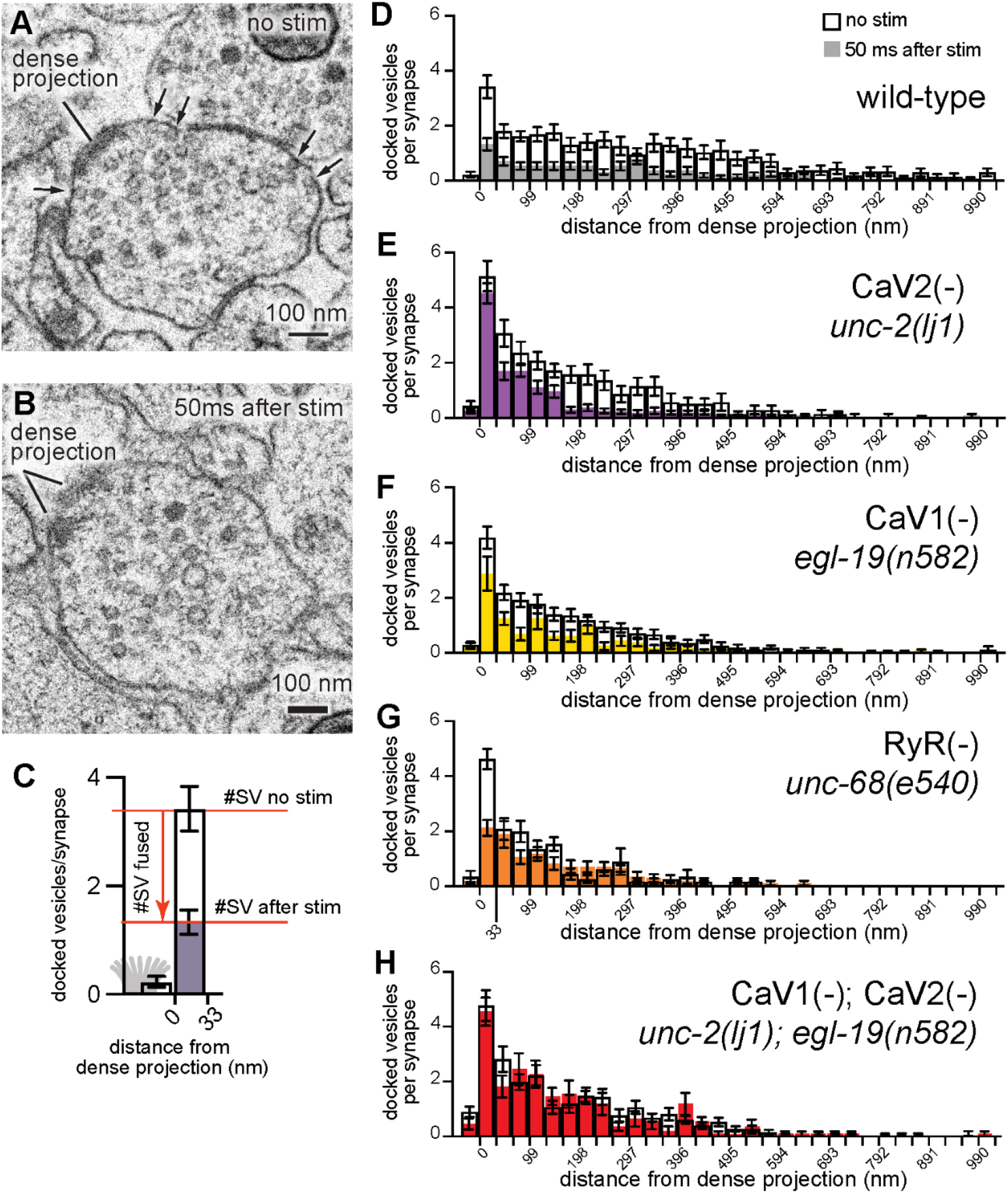
CaV2 and CaV1-RyR act at distinct vesicle release sites. (A-B) Docked vesicles (black arrows) are present near dense projections in electron micrograph of unstimulated animals, but are reduced 50ms after channelrhodopsin stimulation. (C) Interpretation of docking histograms. The number of synaptic vesicles that fuse due to stimulation can be determined by comparing the number of synaptic vesicles docked within the active zone without or with stimulation. (D-H) Average number of docked vesicles per synapse at a given distance from the dense projection with, or without light stimulation of channelrhodopsin. (D) Wild-type animals exhibit fewer docked vesicles at all locations after stimulation. Wild type (no stimulation), n=26 synapses. Wild type (stimulated) n = 24 synapses. (E) The CaV2 null mutant *unc-2(lj1)* fuses vesicles greater than 33 nm from the dense projection but not docked vesicles directly adjacent to the dense projection. CaV2(−) (no stimulation) n=14, CaV2(−) (stimulated) n=27 synapses. (F) The CaV1 hypomorphic mutant *egl-19(n582)* exhibits reduced fusions at all distances. CaV1(−) (no stimulation) n=29 synapses. CaV1(−) (stimulated) n=16 synapses. (G) The RyR mutant *unc-68(e540*) exhibits fusions adjacent to the dense projection, but lacks fusions of lateral vesicles. RyR(−) (no stimulation) n=11 synapses. RyR(−) (stimulated) n=17 synapses. (H) The double mutant CaV1 *egl-19(n582)* and CaV2 *unc-2(lj1)* lack fusion of all docked vesicles after stimulation. CaV2(−) CaV1(−) (control) n=24 synapses. CaV2(−) CaV1(−) (stimulated) n=17 synapses. Errors given in SEM, N = 2 animals for each condition. Micrographs were segmented blind to treatment and genotype. Data available as Source Data 3

To identify vesicle fusions associated with particular calcium channels, we analyzed the distribution of docked vesicles in mutant animals. In unstimulated animals, docked vesicles are clustered around dense projections, although many are observed at lateral regions extending hundreds of nanometers from dense projections. Docked vesicles are uniformly depleted after stimulation in wild-type animals (Figure 3D). Genetic ablation of CaV2 channels reduced vesicle fusions adjacent to the dense projection; docked vesicles distal to the dense projection still fused in response to stimulation (Figure 3E). To assay CaV1 function, we analyzed the hypomorph *egl-19(n582)* to avoid the lethality observed in the double mutants. Mutation of the CaV1 channel reduced fusion broadly, although significant vesicle fusions were observed within 100 nm of the dense projection (Figure 3F). In the absence of RyR only CaV2 is functional, and vesicle fusions are only observed in the 33 nm pool —directly adjacent to the dense projection (Figure 3G). The CaV1 CaV2 double mutant exhibited no change in the number and distribution of docked synaptic vesicles after stimulation (Figure 3H). Together, these data demonstrate that *C. elegans* neuromuscular junctions have two spatially distinct pools of synaptic vesicles: a central pool dependent on CaV2 calcium channels and a lateral pool dependent on CaV1 and RyR.

### CaV2 and CaV1 differentially localized at synapses

The electron microscopy data suggest that CaV1 and CaV2 are localized to spatially separate areas of the active zone. To localize calcium channels with normal expression levels, we tagged the endogenous genes and localized the proteins using fluorescence microscopy. We performed 3-color imaging using dense projection markers as an anatomical fiducial at the center of the synapse. Because *C. elegans* synaptic varicosities are less than 1μm in diameter, super-resolution microscopy was required to resolve channel clusters. A segment of the dorsal nerve cord was imaged, and the region of imaging was restricted to a narrow band to avoid potential complications by CaV1 expression in muscle. To ensure that the pattern of synapses in our fluorescence images matched the arrangement of neuromuscular junctions, we reconstructed 20 µm of the dorsal cord. All imaging was conducted on living, acutely anesthetized nematodes.

Multiple tagging sites were tested for all genes, but in some cases the tags disrupted function or the splice isoforms were not expressed in neurons. Therefore, we tagged internal sites within regions of poor conservation (Extended Data Figure 4A,B). For example, CaV2 was tagged with HALO (Los et al., 2008) in the second extracellular loop near the N-terminus (Kurshan et al., 2018; Schwartz & Jorgensen, 2016).

To confirm that the pattern of calcium channels in our fluorescence images matched the arrangement of dense projections, we reconstructed 20 µm of the dorsal nerve cord from serial sections for electron microscopy (Figure 4A). CaV2 / UNC-2 channels have been localized to the dense projection by immuno-electron microscopy in the nematode (Gracheva, Hadwiger, Nonet, & Richmond, 2008). CaV2 clusters are localized every 1.10 +/- 0.16 µm along the dorsal cord by super-resolution microscopy, similar to the distribution of dense projections in the reconstruction (1.02 / µm). To identify proteins associated with CaV2, we tagged multiple active zone components implicated in scaffolding and release complexes at the dense projection (Ackermann, Waites, & Garner, 2015; Südhof, 2012). Neurexin (*nrx-1*), Magi (*magi-1*), Syde1 (*syd-1*), Liprin-α (*syd-2*), RIMBP (*rimb-1*), and α-Catulin (*ctn-1*) were tagged with SkylanS. Each of these proteins is closely associated with CaV2::HALO puncta placing them at the dense projection as well (Figure 4B).

**Figure 4.**
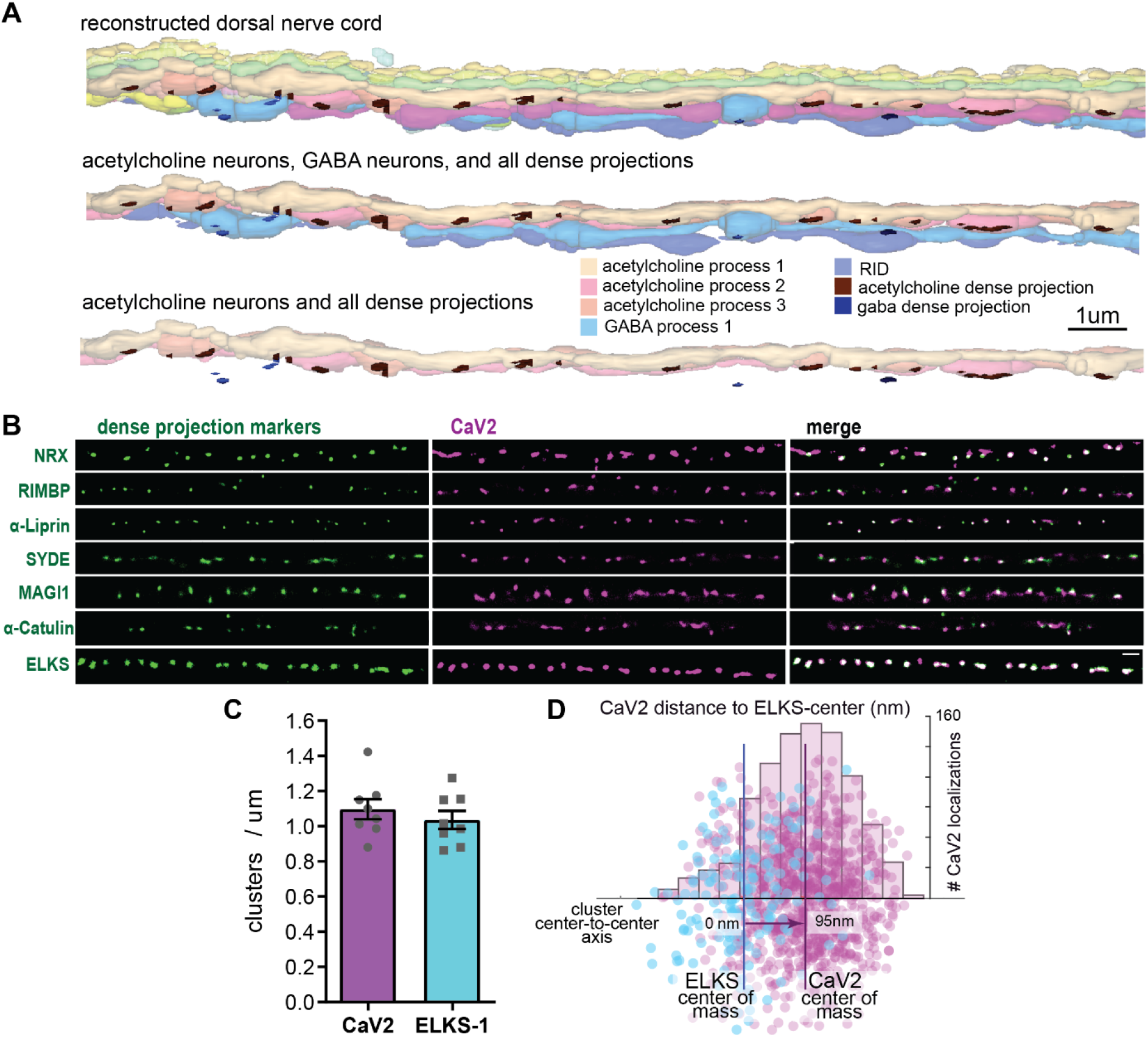
Dorsal nerve cord reconstruction and candidate dense projection markers. (A) 20-micron reconstruction of the wild-type *C. elegans* dorsal nerve cord. Dense projections are highlighted to compare to superresolution images below. Scale bar 1um, section thickness 100 nm. (B) CaV2 colocalizes with cytomatrix active zone proteins. Super-resolution images of Skylan-S-tagged cytomatrix protein homologs in *C. elegans* NRX-1, RIMB-1, SYD-2, SYD-1, MAGI-1, CTN-1, ELKS-1 compared to CaV2-HALO in the same animal. (C) ELKS and CaV2 clusters form approximately 1 / um along the dorsal nerve cord from super-resolution image analysis. Clusters were quantified for over dorsal nerve cords with an average length of 17.8um, N= 8 animals (D) Localization plot tool (Proberuler) example diagram of a single ELKS (cyan) and CaV2 (magenta) synapse. Cluster centers are marked by solid lines. Data available as Source Data 5

ELKS clusters in particular are tightly associated with CaV2 clusters (Figure 4C), and will serve as a synaptic fiducial for CaV2 and CaV1 comparisons. To quantify the distribution of CaV2 relative to ELKS, an axis was plotted between the two cluster centers, and localizations collapsed onto this axis. CaV2 localizations along the axis to the ELKS cluster center were measured and plotted as a histogram (Figure 4D). CaV2 clusters and ELKS clusters are similar in diameter (297 nm vs 294 nm, respectively). However, only 62% of ELKS localizations are within a CaV2 cluster (Figure 5A-C); the cluster centers are slightly offset (124nm). The offset may be because the proteins domains overlap but are not coincident, or alternatively, these proteins may be perfectly colocalized but differ in our plot due to the positions of the tags on the proteins; specifically, CaV2 is tagged on the extracellular side, whereas ELKS is tagged at the C-terminus on the intracellular side. In contrast to the highly concentrated subsynaptic distribution of CaV2, CaV1 is broadly distributed as dispersed puncta (diameter 869 nm), mostly separate from ELKS (262 nm from ELKS center to CaV1 center of mass) (Figure 5A,B), and shares only a 24% overlap with ELKS or CaV2. CaV1 is largely excluded from CaV2 clusters, but the clusters often abut one another (Figure 5C).

**Figure 5.**
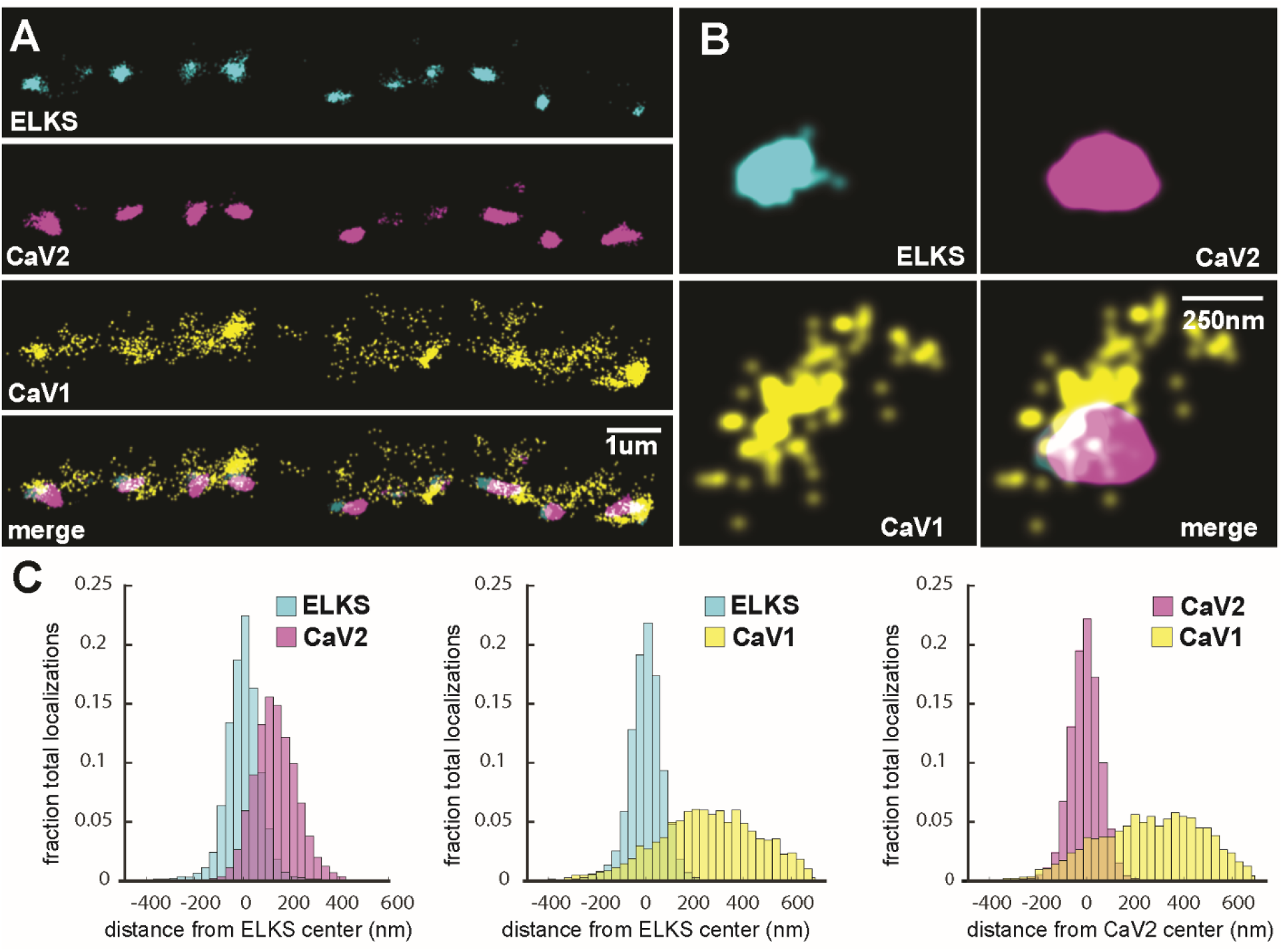
CaV1 is excluded from the dense projection, and dispersed in the active zone. (A, B) Localization microscopy plots of the dorsal nerve cord. ELKS is tagged with Skylan-S. The CaV2-HALO ligand is HTL-JF646, and the CaV1-SNAP tag ligand is STL-JF549pa. (A) CaV2 (magenta) colocalizes with dense projections labeled with ELKS (cyan). CaV1-SNAP (yellow) is largely excluded from the dense projection; and scattered in the synaptic varicosity. Scale bar = 1um. (B) Distributions of CaV2 and CaV1 in a synapse. Dense projections labeled with ELKS (cyan) colocalize with CaV2 (magenta), but not CaV1 (yellow). Scale bar = 250nm. (C) Quantitation of protein localizations from multiple synapses. The center of mass of localizations was calculated from 2D plots. An axis between the centers was fixed and all localizations collapsed onto the axis. Localizations were combined into 33 nm bins, to match the electron microscopy analysis, and plotted as the fraction of total localizations. Data were collected and combined from n=26 synapses, N=5 animals. Data available as Source Data 5

To confirm that the CaV1 localizations are presynaptic and not in the muscle or epidermis, we generated a HALO-tagged CaV1 under the pan-neuronal synaptotagmin promotor (P*snt-1*). This construct was inserted in the CaV1(Δns) strain, and fully rescued CaV1 function (Figure 1B-I). For convenience of genetic crosses, we used RIM binding-protein (RIMBP/RIMB-1) as the dense projection marker. The overexpressed CaV1::HALO tends to be more punctate than the endogenously tagged protein (Extended Data Figure 5A,B). However, CaV1 is not colocalized with RIMBP and the distances between clusters are similar to the endogenously tagged gene (center-to-center CaV1, endogenous tag to ELKS: 262nm; transgene tag to RIMBP: 378 nm)(Extended Data Figure 5C). To demonstrate that CaV1 clusters are presynaptic, we assayed colocalization with a presynaptic marker. LIN7 / VELI is a potential CaV1 scaffolding protein via PDZ domain interactions (Butz, Okamoto, & Südhof, 1998; Pym et al., 2017). We expressed LIN7 in acetylcholine motor neurons using the *unc-129* promotor (Extended Data 5D). CaV1 and LIN7 clusters are closely associated, but not associated with RIMBP (Extended Data Figure 5E). These data suggest that CaV1 is localized at presynaptic boutons in a separate domain from CaV2 channels.

If CaV1 and RyR function in the same vesicle fusion pathway, they should be colocalized (Piggott & Jin, 2021). RyR was tagged with HALO at the N-terminus of the neuronal isoform (Extended Data Figure 4C) (Marques et al., 2020). RyR localizations were compared to CaV1 localizations and a dense projection marker, in this case ELKS-Skylan-S (Figure 6A,B). RyR localizations are diffusely distributed, and lateral to the dense projection (ELKS to RyR center of mass distance: 393 nm; 25 synapses) (Figure 6C). RyR localizations are correlated with CaV1 (RyR to CaV1 center of mass: 166 nm). Closer inspection of the images suggests that RyR and CaV1 are often interdigitated in adjacent zones (Figure 6A). To characterize this relationship, we performed a nearest neighbor analysis: 94% of RyR localizations are within 100nm of a CaV1 localization (Figure 6D). CaV1 exhibits a slightly broader distribution; nevertheless, 82% of CaV1 localizations are within 100nm of a RyR channel. The spatial correlation between CaV1 and RyR is consistent with the functional interactions observed by physiology and electron microscopy at lateral sites, which is independent of CaV2.

**Figure 6.**
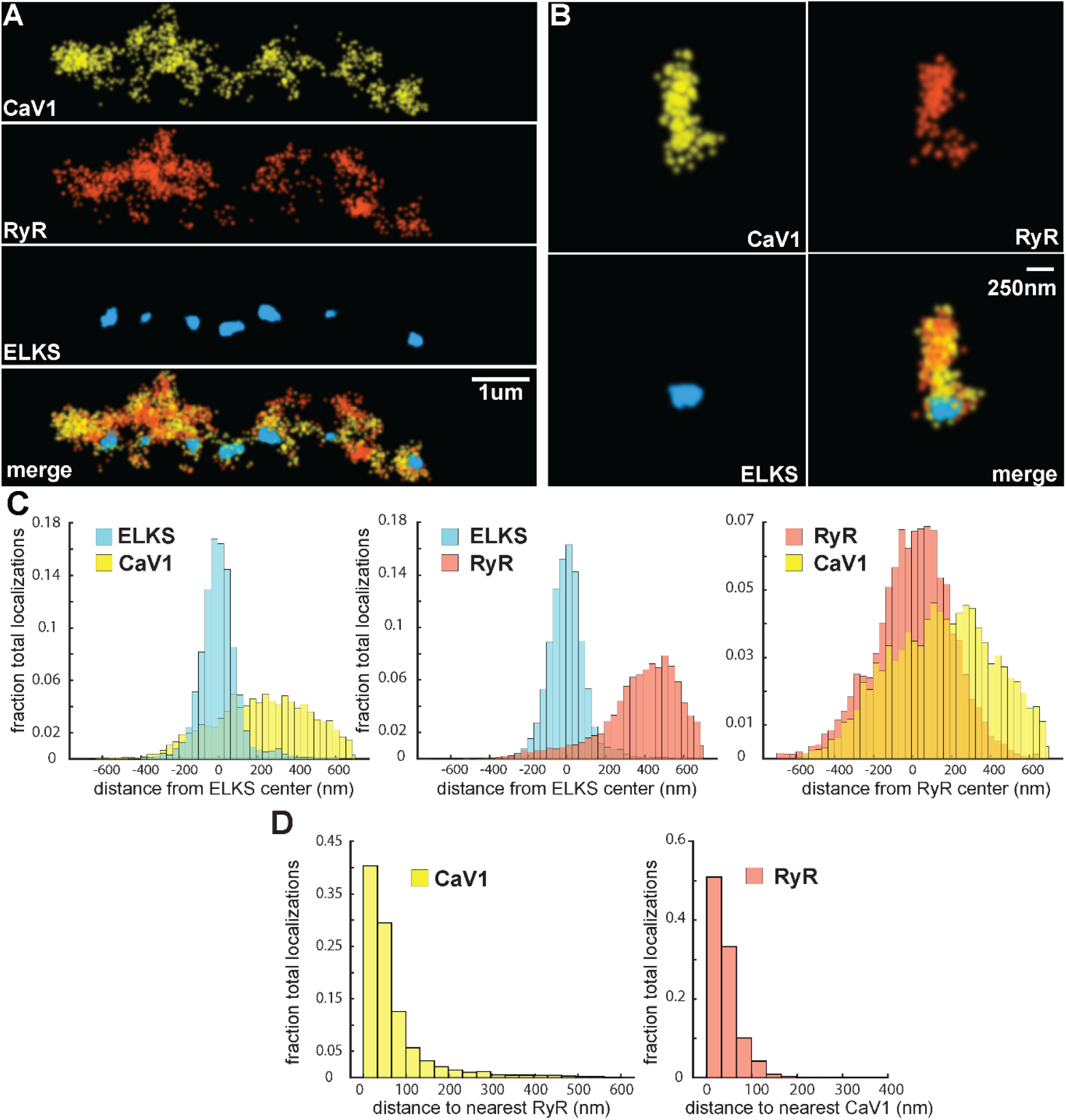
CaV1 and RyR are adjacent. (A) CaV1 and RyR are adjacent along the dorsal nerve cord, lateral to the dense projection. Animals and HTL-JF646. Scale bar = 1um. CaV1-SNAP is labelled with STL-JF549pa, RyR*-*HALO is labelled with HTL-JF646, and dense projections are labeled by ELKS-Skylan-S. (B) RyR and CaV1 colocalize within synapses. Labelling as in ‘A’. Scale bar = 250nm. (C) Distances from CaV1-SNAP localizations to center of ELKS-Skylan-S cluster versus ELKS localizations to ELKS center. Distances from RyR-HALO localizations to center of ELKS-Skylan-S cluster versus ELKS localizations to ELKS center. Distances from CaV1-SNAP localizations to the center of the RyR-HALO cluster versus RyR-HALO localizations to the RyR center. N=5 animals, n=25 synapses. (D) RyR and CaV1 are adjacent. Left, nearest neighbor analysis was performed on CaV1-SNAP localizations to find the nearest RyR-HALO localization. Right, nearest neighbor distances from RyR-HALO to CaV1-SNAP were calculated. n=5 animals, 25 synapses. Data available as Source Data 5

### Different UNC-13 isoforms are associated with CaV1 and CaV2

Vesicle docking and SNARE priming requires UNC-13 proteins. Null mutations in *unc-13* nearly eliminate neurotransmission and vesicle docking in *C. elegans* (Hammarlund et al., 2007; Richmond et al., 1999). We edited the *unc-13* locus at the common C-terminus to append Skylan-S to all isoforms (‘UNC-13all’)(Extended Data 4D). Both CaV2 and CaV1 calcium channels are tightly associated with UNC-13 (Figure 7A-C). Nearest-neighbor analysis indicates that 99.7% of CaV2 channels are within 100 nm of an UNC-13 localization, and 89% of CaV1 channels are within 100 nm of an UNC-13 protein (Figure 7D).

**Figure 7.**
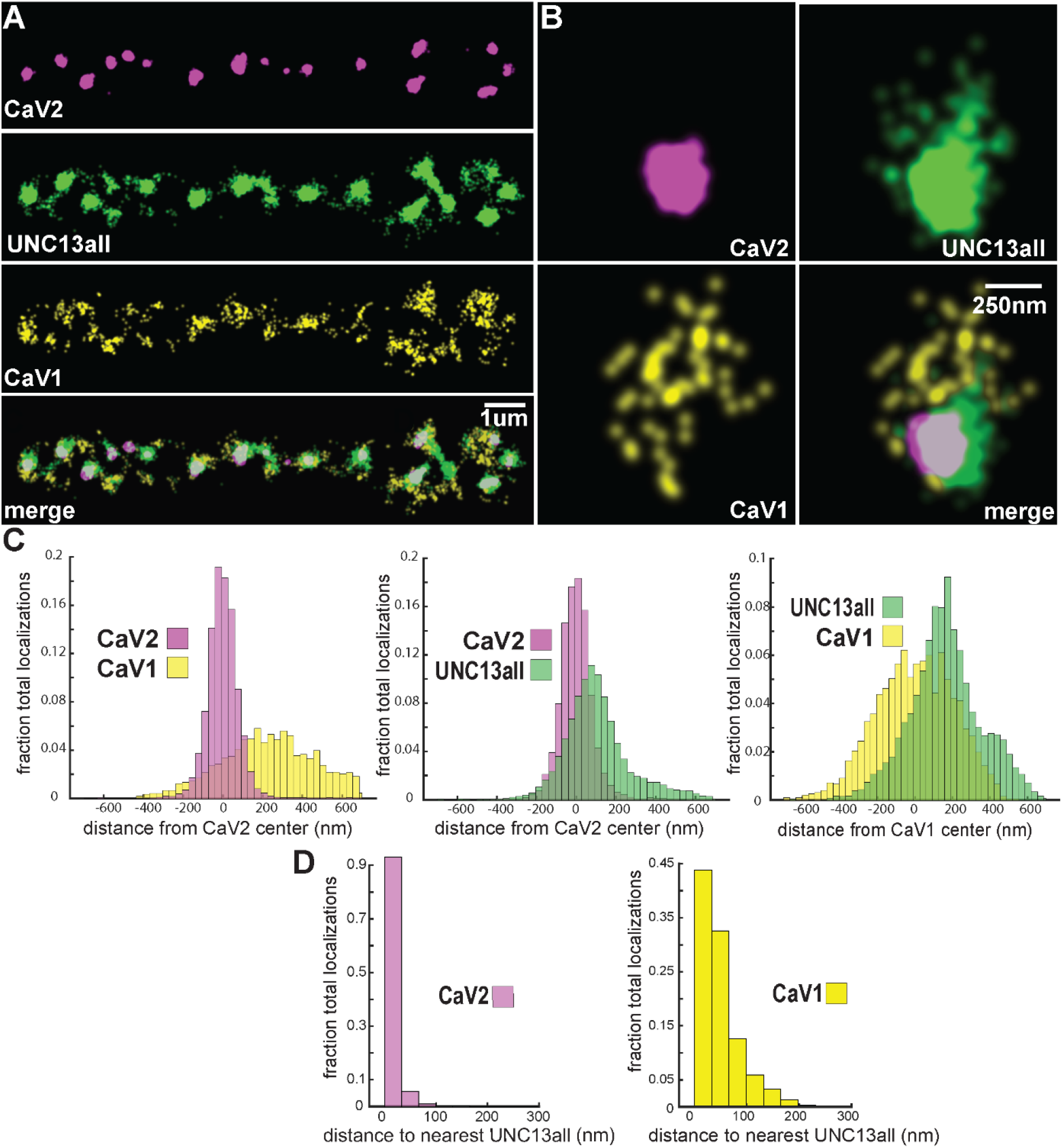
UNC-13 isoforms colocalize with CaV2 and CaV1 calcium channels. Localization microscopy identifies CaV1 and CaV2 associated with ‘UNC13all’, which labels a C-terminal site common to all UNC-13 isoforms. (A) UNC-13all colocalizes with CaV1 and CaV2 along the dorsal nerve cord. Proteins are labelled with CaV2-HALO stained with HTL-JF646, CaV1-SNAP stained with STL-JF549, and UNC13all-Skylan-S. (B) UNC-13all colocalizes with CaV1 and CaV2 within synapses. Staining as in ‘A’. (C) Left, distances from CaV1-SNAP localizations to the center of the CaV2-HALO cluster, and CaV2-HALO localizations to the center of the CaV2-HALO cluster. Middle, distances from UNC13all-Skylan-S localizations to the center of the CaV2-HALO cluster. Right, distances from UNC13all-Skylan-S localizations to the center of the CaV1-SNAP cluster, n=5 animals, 25 synapses. (D) Left, nearest-neighbor distances between UNC13all and CaV1 and CaV2 localizations. Right, nearest neighbor analysis between UNC13all-SkylanS and CaV2-HALO or CaV1-SNAP measured from synaptic regions, n=5 animals, 25 synapses. Data available as Source Data 5

The short isoform UNC-13S lacks the RIM-binding domain C2A, and has a diffuse distribution at synapses (Hu et al., 2013; Weimer et al., 2006; Zhou et al., 2013). UNC-13S was edited at its unique N-terminus to include a Skylan-S tag. UNC-13S does not colocalize with CaV2 (peak-to-peak 319 nm) but is associated with CaV1 (Figure 8A-C). Nearest neighbor analysis indicates that 99% of UNC-13S localizations are within 100nm of a CaV1 channel (Figure 8D), demonstrating that a specialized isoform of UNC-13 docking machinery is localized to CaV1 calcium channels.

**Figure 8.**
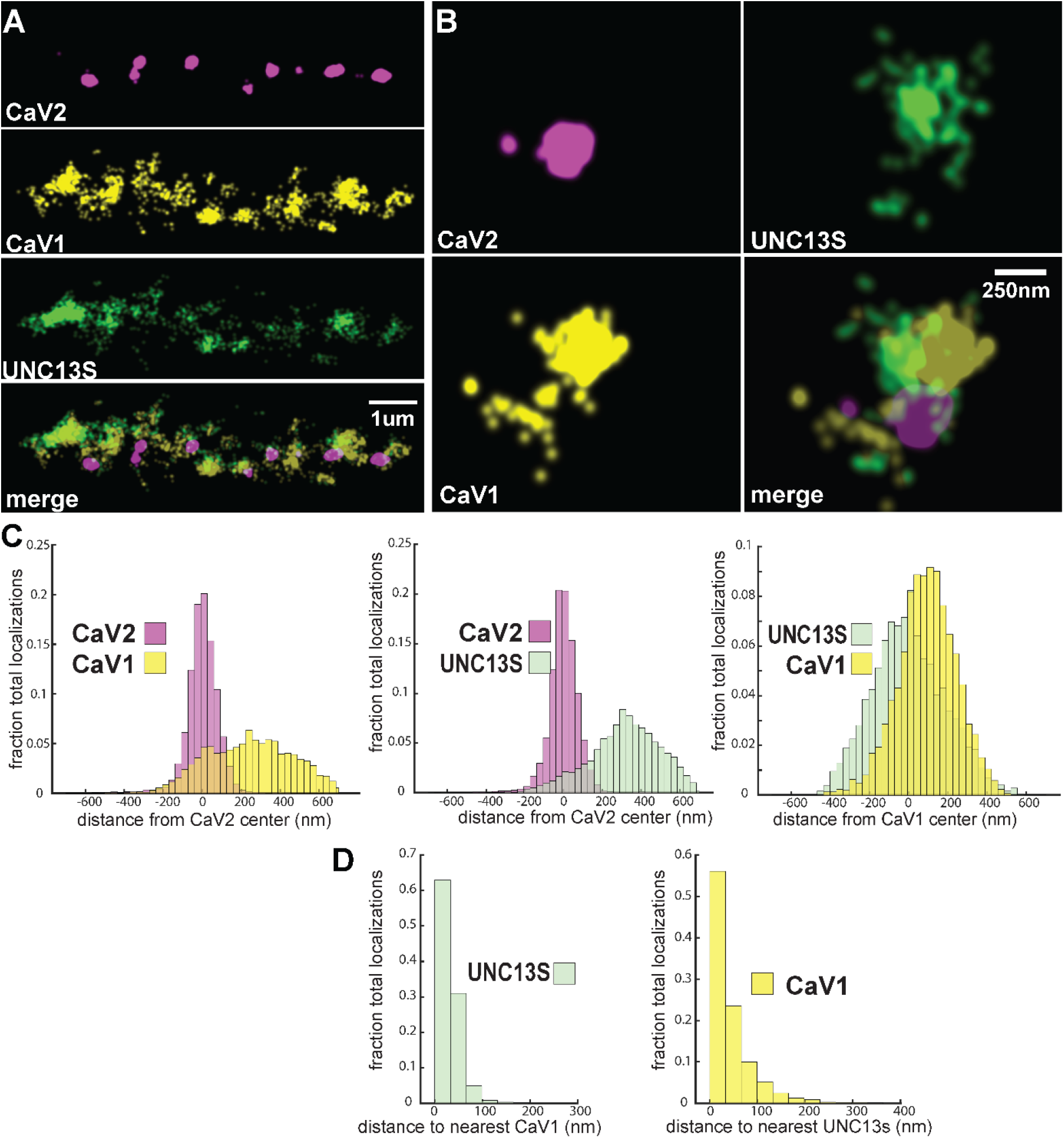
UNC-13S is associated with CaV1 calcium channels. Localization microscopy identifies CaV1 associated with UNC13s which labels a n-terminal site common to a short isoform. (A) UNC-13S localizes with CaV1 along the dorsal cord, but not with CaV2. The endogenous protein tags CaV2-HALO was stained with HTL-JF646, CaV1-SNAP with STL-JF549, and imaged with Skylan-S-UNC13S using single-molecule localization microscopy. (B) UNC-13S localizes with CaV1 within synapses. Strain was labelled as in ‘A’. (C) Left, distances from CaV1-SNAP localizations to the center of the CaV2-HALO cluster compared to CaV2-HALO localizations to their own center. Middle, distances from Skylan-UNC13S localizations to the center of the CaV2-HALO cluster. Right, distances from Skylan-UNC13S localizations to the center of the CaV1-SNAP cluster. N=5 animals, 25 synapses (D) Nearest-neighbor distances of CaV1-SNAP to Skylan-UNC13S localizations. Nearest neighbor analysis of Skylan-UNC13S to CaV1-SNAP measured from synaptic regions. N=5 animals, 25 synapses. Data available as Source Data 5

## Discussion

Calcium channel classes tend to be associated with specific tissue functions: CaV2 (N, P/Q, R-type) with synaptic transmission, and CaV1 (L-type) channels with muscle contraction. Here, we demonstrate that both CaV2 and CaV1 channels drive vesicle fusion at *C. elegans* neuromuscular junctions and mediate the release of different synaptic vesicle pools. In electrophysiological assays, these pools are genetically separable and perfectly complementary. CaV2 channels fuse vesicles near the dense projection, whereas CaV1 channels drive vesicle fusion at lateral sites in the same synapses, as revealed by flash-and-freeze electron microscopy. Super-resolution imaging indicates that CaV2 channels and the active zone proteins Neurexin, α-Liprin, ELKS, and RIMBP are in tight 300 nm clusters at the dense projection, and CaV2 is associated with the long isoform of the docking and priming protein UNC-13L. By contrast, CaV1 is dispersed in the synaptic varicosity and is associated with the short isoform UNC-13S. Finally, vesicle fusion mediated by CaV1 is dependent on the ryanodine receptor, presumably by regulating calcium release from internal stores (Figure 9).

**Figure 9.**
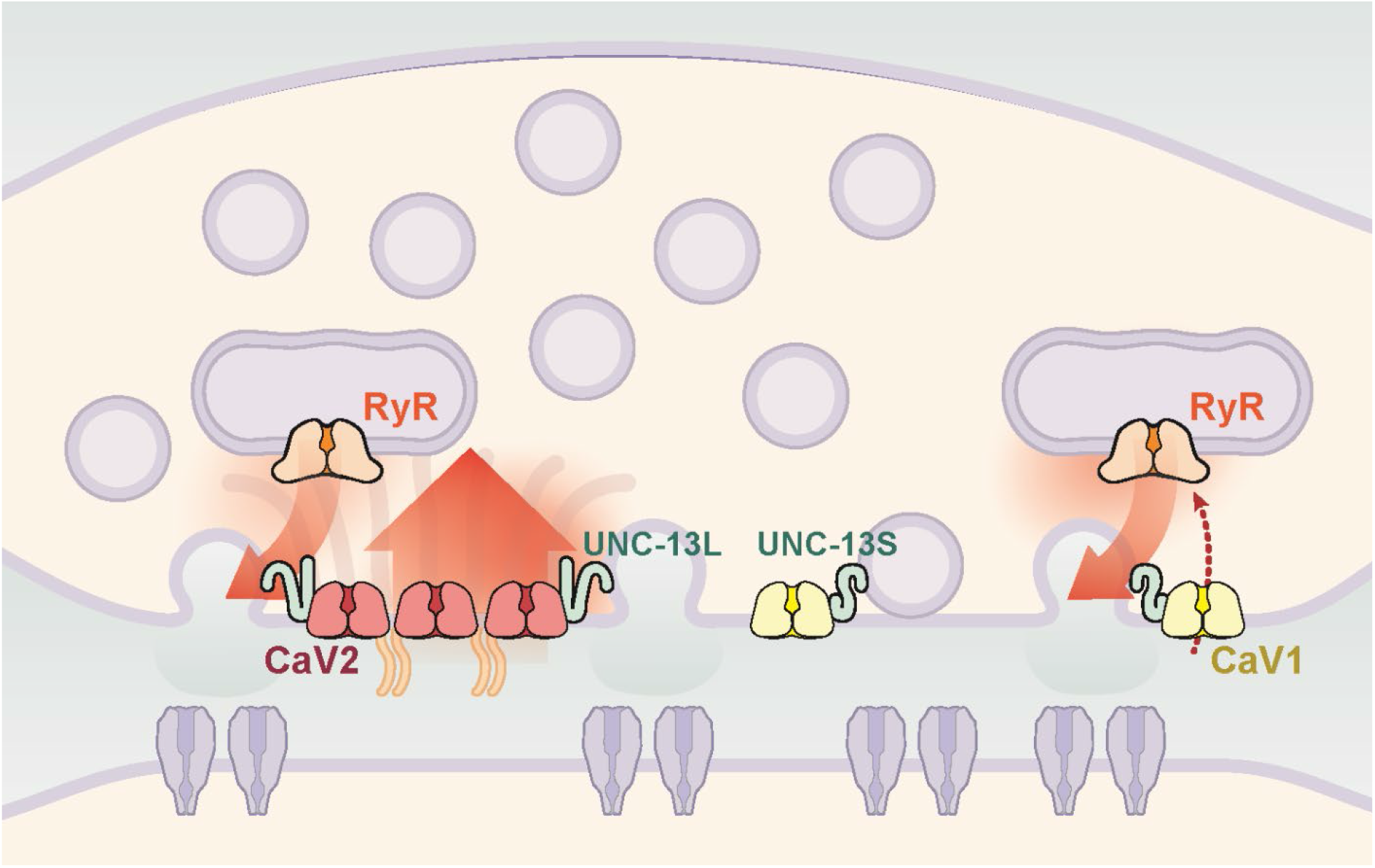
Two independent release sites for synaptic vesicles. Voltage-gated calcium channels localize to two distinct zones at the neuromuscular synapse of *C. elegans*. The CaV2 channel localizes to the dense projection along with ELKS, RIMBP, Neurexin, Liprin-alpha, SYDE, MAGI1, alpha-Catulin and the SNARE priming protein UNC-13L. CaV2 is required to fuse synaptic vesicles are docked directly adjacent to the dense projection. The second channel CaV1 is at a lateral site centered 300nm from the dense projection but can span hundreds of nanometers. CaV1 requires coupling to RyR to synaptic vesicles at the lateral site. These near and far pools utilize specific release machinery. Most UNC-13all localizes to the dense projection. However, some UNC-13 localizes with CaV1 at the lateral site. Isoform specific tagging shows UNC-13S localized with lateral site.

### CaV1 functions at synapses

In *C. elegans* there is only a single L-type channel, encoded by the *egl-19* gene, which primarily plays a role in muscle (Lee et al., 1997). EGL-19 also generates calcium action potentials in some neurons (Liu et al., 2018), and thereby can substitute for the absence of voltage-gated sodium channels in nematodes (Liu et al., 2018). However, L-type channel inhibitors do not affect evoked release in motor neurons, but rather cause a decrease in the mini rate (Tong et al., 2017). We demonstrate that null mutations in *egl-19* have reduced rates of tonic miniature currents, are localized presynaptically, and are involved in fusions of a specific sub-pool of synaptic vesicles. Together these data demonstrate that this L-type channel is acting to mediate fusion of synaptic vesicles.

### Multiple calcium channels - coupling and voltage-dependence

Based on physiological data alone, it was possible that CaV1 and CaV2 channels could function at separate synapses. However, the electron microscopy and fluorescence experiments demonstrate that these channels are localized and functioning at the same synaptic varicosity. Participation of multiple classes of calcium channels at the same synapse may function to tune the dynamics of neurotransmission. Differences in voltage-dependent activation, inactivation, clustering, or distance to docked vesicles could regulate synchronous or asynchronous release (Dolphin, 2021).

One important difference is that CaV1 channels inactivate more slowly than CaV2 channels and can therefore more accurately report synaptic depolarization (Naranjo, Wen, & Brehm, 2015; Yu, Yuan, Westenbroek, & Catterall, 2018). For example, CaV1.3 and CaV1.4 mediate vesicle fusion in sensory neurons, and slow inactivation of these channels allows these synapses to accurately report the depolarization status of the synaptic bouton (McRory et al., 2004; Platzer et al., 2000). The *C. elegans* CaV1 channel EGL-19 also exhibits slow inactivation *in vivo* (Lainé, Ségor, Zhan, Bessereau, & Jospin, 2014), which likely contributes to tonic miniature currents in a graded fashion (Liu et al., 2018). Continued calcium influx through CaV1 may act to terminate neurotransmission in motor neurons; calcium influx through EGL-19 is specifically coupled to repolarization via SLO-2 BK potassium channels (Liu et al., 2014).

The most profound difference for CaV1 and CaV2 channels at *C. elegans* neuromuscular junctions is that they are differentially localized: CaV2 is localized to a large cluster at the dense projection, CaV1 is distributed broadly in the synapse. Vesicle pools can be assayed as tightly coupled or loosely coupled to calcium channels based on sensitivity to EGTA (Dittman & Ryan, 2019; Eggermann et al., 2012). At *C. elegans* neuromuscular junctions, UNC-13L mediates tight coupling (EGTA-insensitive), whereas UNC-13S mediates loose coupling (EGTA-sensitive) (Hu et al., 2013). Consistent with these findings, UNC-13L is tightly coupled to CaV2 channels at dense projections. CaV1 channels are dispersed across the synapse and localizations are frequently solitary. Nevertheless, CaV1 channels can drive fusion across a broad distribution of docked vesicles, extending 500 nm from the dense projection.

The requirement of CaV1 for the fusion of this distal pool of vesicles is likely to be mediated by the ryanodine receptor rather than by CaV1 directly. Calcium influx from CaV1 channels stimulates the release of calcium from the endoplasmic reticulum via the ryanodine receptor (Bouchard, Pattarini, & Geiger, 2003). In skeletal muscle, CaV1.1 is physically coupled to RyR1 and voltage-sensing by the calcium channel can gate the ryanodine receptor in the absence of extracellular calcium (Schneider, 1994). In neurons, it is not clear if ryanodine receptor activation is physically coupled to CaV1 activation. In *C. elegans*, there is likely no direct-physical link between CaV1 and RyR since depolarizations in the absence of calcium do not elicit synaptic vesicle release (Liu et al., 2005). Nevertheless, our data indicate that these channels are linked spatially and functionally. Nearest neighbor analysis indicates that essentially all RyR localizations are within 100 nm of a CaV1 channel, and the electrophysiology and electron microscopy demonstrate that they mediate fusion of the same pool of vesicles.

In contrast to CaV1, CaV2-mediated release is not dependent on RyR but rather is potentiated by RyR. In the presence of CaV1 blockers, the frequency of miniature currents is not decreased by loss of the RyR, indicating that CaV2 reliably drives vesicle fusion on its own. Rather, it is the amplitude of the miniature current that is reduced in the absence of RyR. The simplest interpretation is that calcium sparks from internal stores are directly acting to fuse synaptic vesicles (Llano et al., 2000); however, we cannot exclude that potentiation is caused indirectly by increases in basal cytosolic calcium at synapses. It is possible that increases in global calcium are acting to increase the probability of vesicle fusion (Galante & Marty, 2003).

The participation of the CaV1 L-type channel EGL-19 in synaptic transmission in *C. elegans* is unusual but not unprecedented. CaV1 channels also play a primary role in synaptic vesicle fusion at graded sensory synapses: CaV1.3 drives neurotransmission in hair cells and CaV1.4 acts at ribbon synapses in photoreceptors (Schmitz & Witkovsky, 1997; Zhang, Robertson, Yates, & Everett, 1999). CaV1.2 and CaV1.3 are also expressed broadly in the brain and function in dendritic spines during synaptic plasticity (Hell et al., 1996, 1993; Nanou & Catterall, 2018). Ryanodine receptors are also found at vertebrate presynapses (Bouchard et al., 2003), and potentiate release at GABA neurons in the cerebellum (Galante & Marty, 2003). CaV1.3 is physically coupled to and activates RyR2 to release calcium stores in cultured hippocampal neurons (Kim et al., 2007). Recent work indicates that CaV1 channels and RyR2 must be colocalized to function together at cellular junctions in the cell body of hippocampal neurons (Sahu et al., 2019).

Moreover, cooperation between CaV2 and CaV1 channels at synapses may be widespread. Immunofluorescence experiments indicate that both CaV1 and CaV2 channels are localized together at the same neuromuscular junctions in the fly (Krick et al., 2021), and mouse CaV1 and CaV2 channels function together at neuromuscular junctions (Katz, Ferro, Weisz, & Uchitel, 1996; Urbano & Uchitel, 1999). Pharmacological experiments suggest that CaV1 and CaV2 channels may function together in GABA neurons in the central nervous system (Goswami, Bucurenciu, & Jonas, 2012; Rey et al., 2020). Vertebrate homologs of the two worm UNC-13 isoforms, Munc13-1 and bMunc13-2, function together at the same synapses (Kawabe et al., 2017) and suggests that this organization of synapses into separately regulated pools of synaptic vesicles may be general.

## Methods

### Rescue of Lethal Calcium Channel Mutants

Lethal CaV1 / *egl-19(st556)* animals were rescued by Mos-mediated transgenes (*oxTi1047[Pset-18::egl-19b::let-858 3’utr] II*. EG9034 ‘CaV1 (Δns)’) or by extrachromosomal array (*oxEx2017[Pset-18::eGFP_egl-19b::let858utr ; Punc-122::GFP].* EG8827 ‘CaV1(Δns) RyR(−)’) (Frøkjær-Jensen et al., 2014, 2008). An *egl-19* minigene was constructed from cDNA and portions of gDNA containing small introns to aid expression. The first exons 1-4 are cDNA, followed by gDNA of exon 5-9, and cDNA of exon 10-17. The minigene was placed downstream from a muscle P*set-18* promoter and inserted directly into the genome by MosSCI (Frøkjær-Jensen et al., 2008).

For the array rescue of CaV1 in muscle, *Pset-18::eGFP_egl-19b::let858utr ; Punc-122::GFP* was microinjected into the gonad of adult hermaphrodite *egl-19(n582) C. elegans.* Array positive animals were selected and crossed with *egl-19(st556)* (RW3563), which rescued lethality but lacked expression in the nervous system (EG8409). The resulting construct *oxTi1047* was crossed into CaV1(−) / *egl-19(st556)* animals (RW3563), which rescued lethality but lacked expression in the nervous system.

To demonstrate that phenotypes in this EG9034/EG8409 were due to loss of nervous system function, we expressed the *egl-19* minigene under the neuron-specific P*snt-1* promoter and inserted the construct into the genome by miniMos (Frøkjær-Jensen et al., 2014). The resulting *oxTi1049* construct was crossed into the muscle-rescued CaV1(Δns) animals (EG9034) to generate EG9145.

Lethal double mutants of CaV2-RyR (genotype: *unc-2 (lj1); unc-68 (e540)*) and CaV2-CaV1 (genotype: *unc-2 (lj1); egl-19 (st556)*) were rescued by an extrachromosomal array expressing SNAP::CaV2/*unc-2* cDNA in a minimum set of acetylcholine head neurons, using a previously described truncated *unc-17* promoter, referred to as ‘P*unc-17*h’ (Hammarlund et al., 2007; Topalidou et al., 2016). The extrachromosomal array *oxEx2096* was generated in the *unc-2(lj1)* strain AQ130 and crossed to RyR / *unc-68(e540)* or CaV1 / *egl-19(st556) oxTi1047[*P*set-18::egl-19b]* animals to generate double mutants. The resulting strains are lethal without the presence of *oxEx2096[*P*unc-17h*::*SNAP*::*unc-2]* and were used in electrophysiology experiments.

### Behavioral Experiments

Animals were maintained under standard laboratory conditions. For behavioral experiments, 3 to 6 well-fed, young adult worms were transferred to a 10cm containing standard NGM. Each assay was recorded for 5 minutes at 8 frames per second using the worm tracking software WormLab (2019.1.1, MBF Bioscience). The trajectory of each worm was collected using WormLab and imported into custom written R scripts for analysis. Worms that crawled out of the field of view during the first 3 minutes were discarded from analysis. Worms whose speed was lower than 100um/s were excluded as they may have been damaged during transfer, the number of worms that fell in this category were few and not different between groups. A reversal was defined as backwards locomotion that lasted more than 4 frames or 500ms.

### Generation of CaV2::HALO by CRISPR/cas9

CaV2 was tagged by CRISPR-mediated insertion of HALO coding DNA into the *unc-2* endogenous genomic locus. A DNA mix containing 1) PCR-generated DNA repair template that includes the HALO tag with an embedded *Cbr-unc-119(+)* cassette flanked by loxP sites and 33bp homology arms to the cut site, 2) plasmid DNA that directs expression of Cas9 and an sgRNA (Schwartz & Jorgensen, 2016), and 3) an inducible negative selection plasmid directing expression of a histamine-gated chloride channel in neurons, pNP403 (Pokala, Liu, Gordus, & Bargmann, 2014) was injected into the gonads of young adult EG6207 *unc-119(ed3)* animals (Maduro & Pilgrim, 1995; Schwartz & Jorgensen, 2016; Zhang et al., 2015). Transgenic animals were selected for expression of *unc-119(+)*, and extrachromosomal-array bearing animals were selected against by addition of histamine to the media. The *loxP::Cbr-unc-119(+)::loxP* region of the insertion was excised by injecting pDD104[Peft-3::Cre] and identifying *unc-119(−)* animals (Dickinson, Ward, Reiner, & Goldstein, 2013). The modified locus introduces HALO-tag within an unconserved region in the second extracellular loop of CaV2 encoding UNC-2a. The resulting strain EG9823 (genotype: *unc-119*(ed3); *unc-2*(ox672[HALO])) was subsequently used to generate CRISPR-mediated insertions of SkylanS tags.

### Generation of Super-Resolution Tags by CRISPR/cas9

Tags for other genes, including *egl-19, unc-68, elks-1, nrx-1, rimb-1, elks-1, syd-2, syd-1, magi-1, ctn-1, unc-13,* and *unc-13b* were constructed as previously described (Schwartz & Jorgensen, 2016). A single plasmid containing sgRNA and the repair template, composed of 57bp homology arms and SkylanS (Zhang et al., 2015) containing a *loxP::Cbr-unc-119(+)::loxP*, was appended by SapTrap plasmid assembly. Each assembled plasmid was mixed with plasmids to express Cas9 in the germline, and HisCl- in neurons, and injected into the gonads of young adult EG9823 animals. After selecting for *unc-119(+)* and selecting against extrachromosomal arrays by histamine application, animals were injected with pDD104[P*eft-3::Cre*], selected for excision of *loxP::Cbr-unc-119(+)::loxP*, and outcrossed once before analysis by super-resolution microscopy.

### Strains

All strains were maintained at 22°C on standard NGM media seeded with OP50.

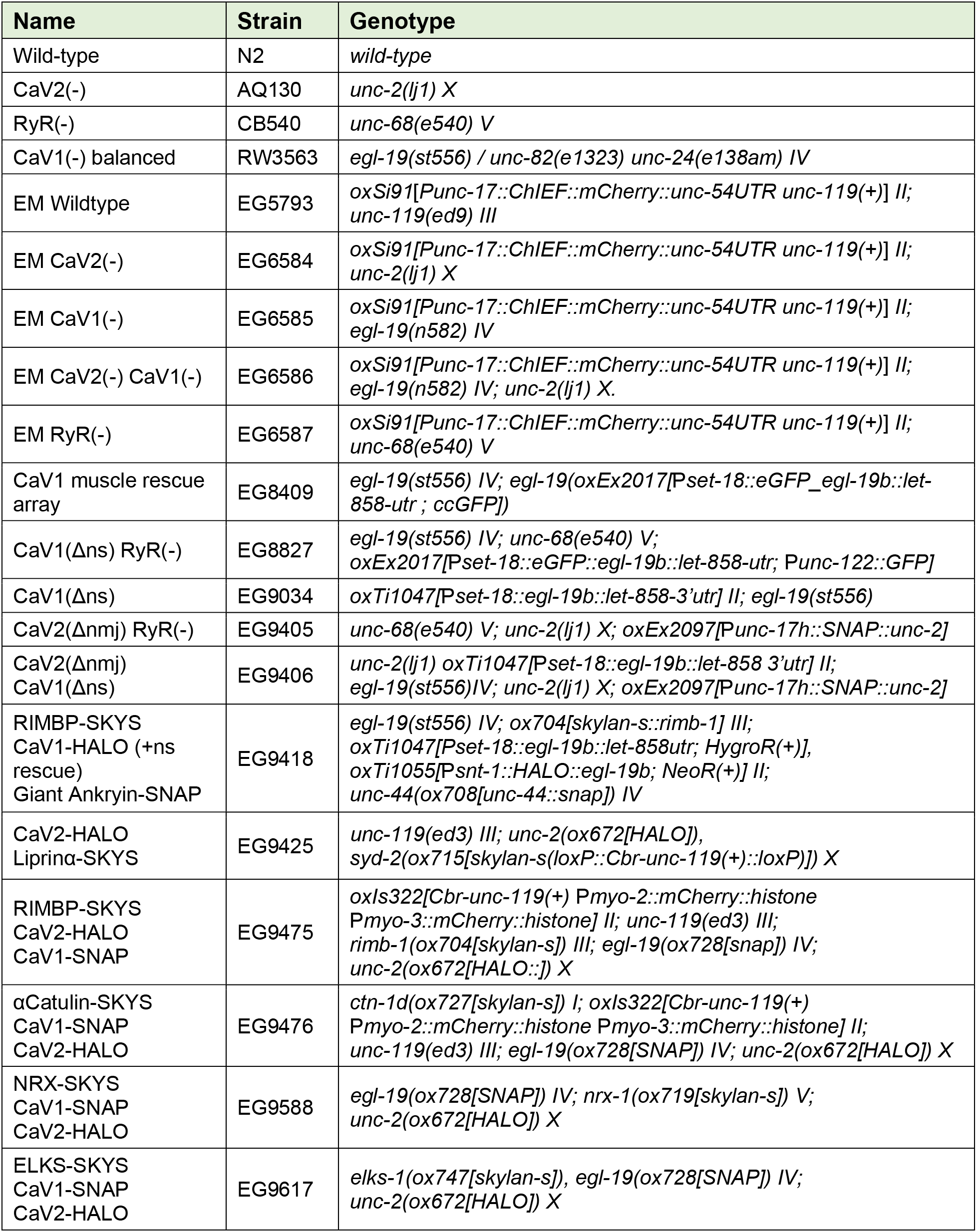
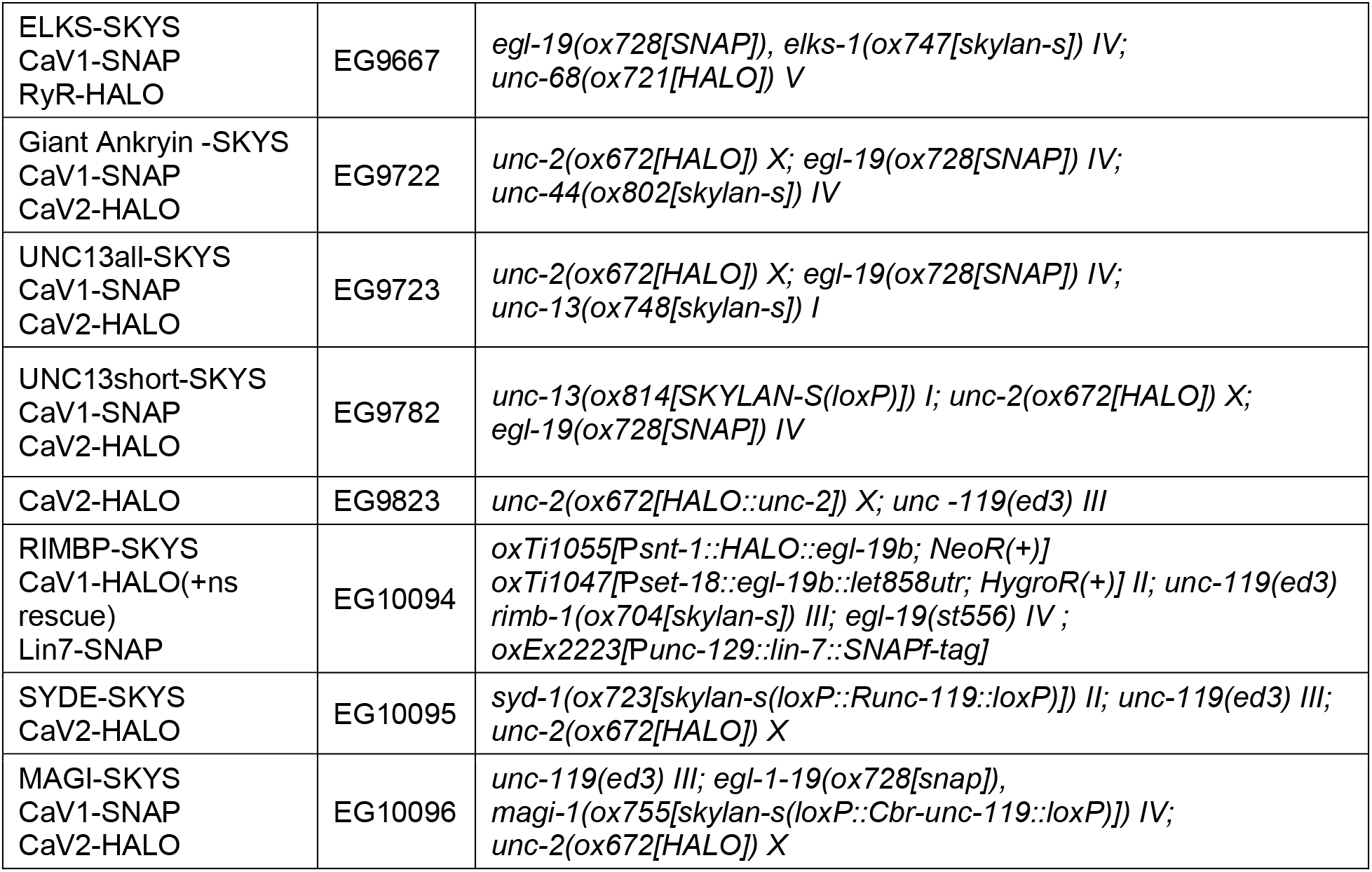

### Single Molecule Localization Microscopy

Super-resolution images were recorded with a Vutara SR 352 (Bruker Nanosurfaces, Inc., Madison, WI) commercial microscope based on single molecule localization biplane technology (Juette et al., 2008; Mlodzianoski, Juette, Beane, & Bewersdorf, 2009). *C. elegans* expressing HALO-tagged proteins (Encell, 2013; Mollwitz et al., 2012) were stained for two hours in 50μM of HTL-JF646, and 50μM of STL-JF549cp, STL-JF549, or STL-JF549pa (Gift from Luke Lavis, Janelia Farms; (Grimm et al., 2017, 2015)). Early super-resolution experiments were conducted with JF549-STL or PA-JF549-STL, we later found that a new cell permeable variant cp-JF549-STL improved labeling of channels. Animals were recovered 12 hours at 15degC on agar seeded with OP50 bacteria. Live intact animals were anesthetized in 25mM NaN3 and regions of their dorsal cords that were positioned directly against the cover glass and away from the intestine were imaged with 640nm excitation power of 10kW/cm2, or 549nm excitation power of 5kW/cm2 SkylanS was imaged by 488nm excitation at 2kW/cm2, while photoactivated by 0.37mW/cm2 405nm light. Images were recorded using a 60x/1.2 NA Olympus water immersion objective and Hamamatsu Flash4 V1 sCMOS, or 60x/1.3 NA Silicon immersion objective and Orca Fusion BT SCMOS camera with gain set at 50 and frame rate at 50 Hz. Data was analyzed by the Vutara SRX software (version 7.0.0rc39). Single molecules were identified by their brightness frame by frame after removing the background. Identified molecules were localized in three dimensions by fitting the raw data in a 12×12-pixel region of interest centered around each particle in each plane with a 3D model function that was obtained from recorded bead data sets. Fit results were filtered by a density based denoising algorithm to remove isolated particles. The experimentally achieved image resolution of 40nm laterally (x,y) and 70 nm axially (in z) was determined by Fourier ring correlation. Localizations were rendered as 80nm.

### SML Analysis

Localization data was exclusively collected from the dorsal nerve cord, which contains axons and synapses but no neuronal soma. We performed a 3D reconstruction of C. elegans dorsal nerve cord to inform region of interest selection from fluorescent images. The orientation of dorsal cord synapses is predictable. Excitatory acetylcholine neurons and inhibitory GABA neurons synapse onto muscle arms (Figure 3A). These connections are near the edges of the cord bundle. Thus, the roll of the animal affects the orientation of the synapse; en face or axial. For single molecule localization experiments, animals were rolled to ensure en face orientation of synapses. Synapses that were in focus and en face were analyzed. The average size of a synapse from the dorsal nerve cord is 579.7nm (SEM +/- 16nm). Thus, super-resolution analysis regions of interest were narrowed to localizations within 700nm of the dense projection marker. Localization position data was flattened in the z-dimension due to chromatic aberrations. A script was used to calculate the center of each probe. To compare the distribution of probe A to probe B, an angle between the two clusters centers was calculated. The distribution distances were calculated by measuring the distance along the center-to-center axis from a probe B to the center of cluster A, and cluster B. Nearest neighbor analysis was done with knnsearch(). The 95% confidence interval of these distance measurements is considered the diameter of the cluster. Distribution center and range or “diameter” were reported as (mean, 95%CI). Proberuler available at https://github.com/bdmscience/proberuler

### Electrophysiology

All electrophysiological experiments were completed with young adult hermaphrodites. The animals were immobilized and dissected as previously described (Ping Liu, Chen, Mailler, & Wang, 2017). Worm preparation was bathed with an extracellular solution containing (in mM) NaCl 140, KCl 5, CaCl2 0.5, MgCl2 5, dextrose 11 and HEPES 5 (pH 7.2). Spontaneous postsynaptic currents (PSCs) at neuromuscular junction were recorded at a holding voltage of -- 60 mV with a pipette solution containing (in mM) KCl 120, KOH 20, Tris 5, CaCl2 0.25, MgCl2 4, sucrose 36, EGTA 5, and Na2ATP 4 (pH 7.2). The classic whole-cell recordings were performed with a Multiclamp 700B amplifier (Molecular Devices, Sunnyvale, CA, USA) and the Clampex software (version 10, Molecular Devices). Data were filtered at 2 kHz and sampled at 10 kHz. Nemadipine-A (Sigma-Aldrich) was first dissolved in DMSO to make frozen stock solution (10mM), and was diluted to a final concentration of 10 µM in extracellular solution before use. Animals were treated for 5 minutes. The frequency and amplitude of minis were quantified with MiniAnalysis (Synaptosoft, Decatur, GA, USA). The amplitudes of evoked currents were quantified using Clampfit (version 10, Molecular Devices, Sunnyvale, CA, USA)

### Quantal Modelling

A 5-term gaussian distribution was fit to cumulative frequency distribution of amplitudes per condition in 1pA bins using MATLAB Curve Fitter (Mathworks, Natick, MA, USA). The terms were centered on 9 +/- 2 pA intervals which represent the mode of amplitudes; a single quantum. (del Castillo & Katz, 1954). The coefficient for the mean of the first gaussian curve was set to the mode amplitude (9 pA+-2). Every coefficient for subsequent gaussian terms were set to 9pA+-2 intervals, but the other coefficients were not constrained and allowed to find a best fit. Area under each curve was calculated using MATLAB trapz().

### Flash and Freeze Electron Microscopy

Electron microscopy was performed as previously described (Watanabe et al., 2013). Freezing was performed on a Leica EMpact2 (Leica, Wetzlar, Germany). To stimulate neurotransmission animals were exposed to blue (488nm) LED light for 20ms and frozen 50ms later. 33nm serial sections were taken and imaged using a Hitachi H-7100 transmission electron microscope equipped with a Gatan Orius digital camera (Gatan, Pleasanton, CA). Micrographs were analyzed in ImageJ using a program for morphological analysis of synapses (Watanabe, Davis, Kusick, Iwasa, & Jorgensen, 2020). Scripts available at: https://github.com/shigekiwatanabe/SynapsEM

### Dorsal Nerve Cord Reconstruction

Serial sections were cut at 100nm and imaged using JEOL JEM-1400 (JEOL, Peabody, MA) then annotated and assembled using TrackEM2 in FIJI (Cardona et al., 2012). Specifically, a wireframe was fit through each process that was suspected to be in the previous micrograph. Then an outline of the plasma membrane of each process was drawn. We analyzed several criteria to more specifically determine the specific process name and type: the morphology of each process and compared to previously published data (J.G. White, E. Southgate, J.N. Thomson, & S. Brenner, 1986), and the number of synapses. These data allow us to determine the identity of a process with some certainty.

**Table 1:**
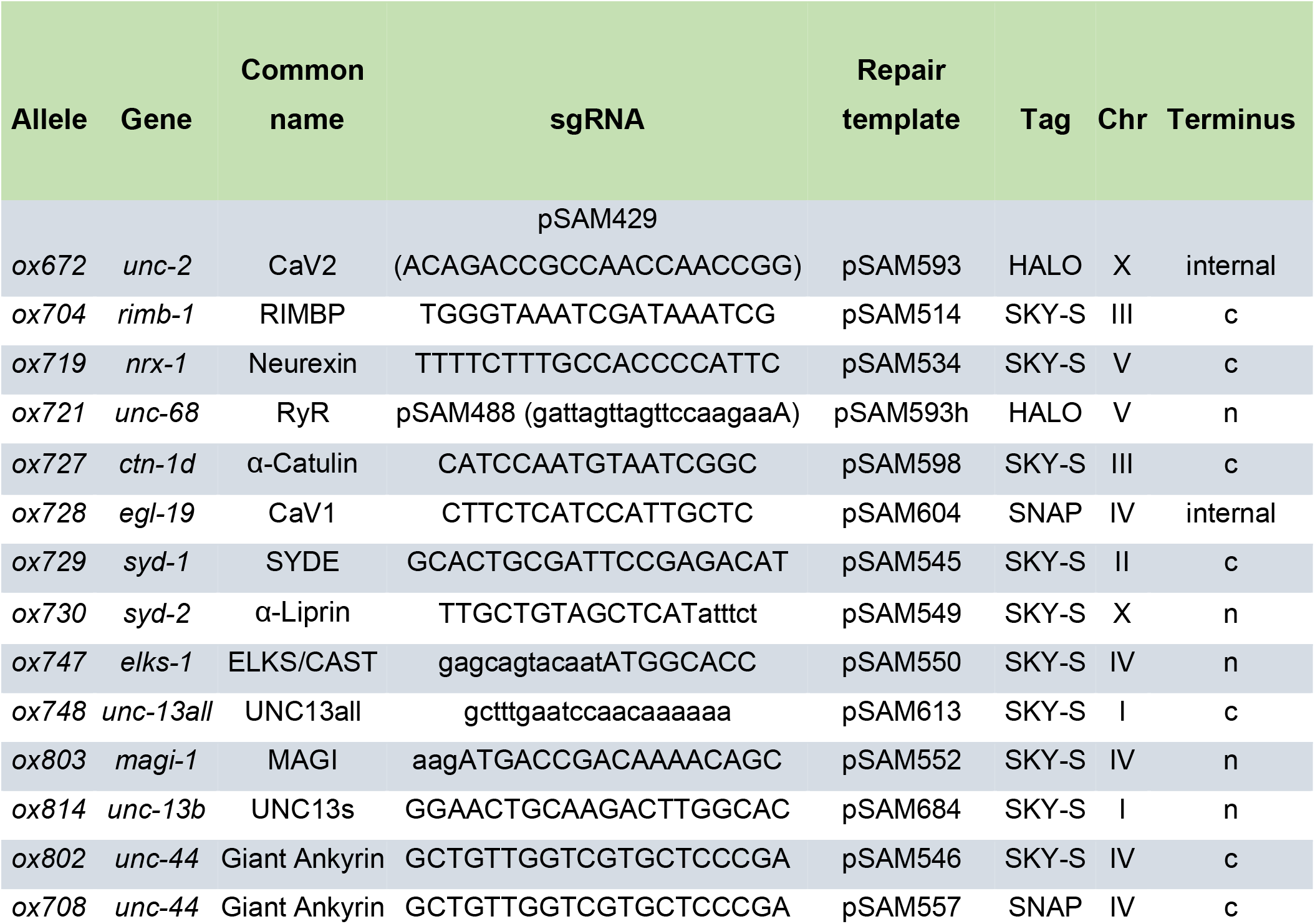
Super-resolution Alleles Generated for this study.

**Table 2:** Common names and nomenclature used in this study.

| Common name | Mammalian ortholog | *C. elegans* ortholog | *Drosophila* ortholog |
| --- | --- | --- | --- |
| CaV2 | CaV2.1<br>CaV2.2<br>CaV2.3 | UNC-2 | Cacophony |
| CaV1/L-type | CaV1.1<br>CaV1.2<br>CaV1.3<br>CaV1.4 | EGL-19 | DmCa1D |
| ELKS | ELKS/CAST | ELKS-1 | Bruchpilot |
| RyR | RYR1<br>RYR2<br>RYR3 | UNC-68 | dRyr |
| UNC13all | Munc13-1<br>ubMunc13-2<br>bMunc13-2<br>Munc13-3 | UNC-13L,<br>UNC-13S | Unc13A<br>Unc13B |
| UNC-13S | bMunc13-2<br>Munc13-3 | UNC-13S | UNC13B |
| RIMBP | RIMBP | RIMB-1 | Rbp |
| Veli/LIN7 | LIN7A | LIN-7 | Lin-7 |
| Giant Ankyrin | gAnkB | UNC-44L | AnkG |

| Neurexin/NRX | Neurexin 1 | NRX-1 | DNrx |
| --- | --- | --- | --- |
| α-Liprin | α-Liprin | SYD-2 | Liprin-α |
| SYDE | SYDE | SYD-1 | Syd-1 |
| α-Catulin | α-Catulin | CTN-1 | α-Cat |
| MAGI1 | MAGI1 | MAGI-1 | Magi |

**Table 3:** Summary of strain nomenclature and alleles.

| Common name | Usage | CaV2 | CaV1 | RyR | CaV2 rescue | CaV1 rescue |
| --- | --- | --- | --- | --- | --- | --- |
| Wildtype<br>N2 Bristol | ePhys<br>EM<br>behavior | wt | wt | wt | n/a | n/a |
| CaV1(rescue) | behavior<br>SMLM | wt | *st556* | wt | n/a | *oxTi1047[Pset-18::egl-19b]*<br>*oxTi1049[Psnt-1::HALO::egl-19b]* |
| CaV1(Δns) | behavior<br>ePhys | wt | *st556* | wt | n/a | *oxTi1047[Pset-18::egl-19b]* |
| CaV1(Δns)<br>RyR(-) | ePhys | wt | *st556* | *e540* | n/a | *oxEx2017[Pset-18::eGFP::egl-19b]* |
| CaV2(Δnmj)<br>CaV1(Δns) | ePhys | *lj1* | *st556* | wt | *oxEx2096[Punc-17h::SNAP::unc-2]* | *oxTi1047[Pset-18::egl-19b]* |
| CaV2(Δnmj)<br>RyR(-) | ePhys | *lj1* | wt | *e540* | *oxEx2096[Punc-17h::SNAP::unc-2]* | n/a |
| CaV2(-) | ePhys<br>EM | *lj1* | wt | wt | no | n/a |
| RyR(-) | ePhys<br>EM | wt | wt | *e540* | n/a | n/a |
| CaV1(-)<br>CaV2(-) | EM | *lj1* | *n582* | wt | no | no |
| CaV1(-) | EM | wt | *n582* | wt | n/a | no |

## Acknowledgements

We thank Lexy von Diezmann for development of Proberuler. We thank Luke Lavis for providing all Janelia Fluor dyes (JF Dyes). Wayne Davis suggested using the early expressing *set-18* promoter to rescue CaV1. We thank the University of Utah Fluorescence Imaging Core for instrumentation, and the Caenorhabditis Genetics Center (CGC) for maintaining and distributing strains to the *C. elegans* community. We thank Patrick McEachern, Matthew Rich, Jessica Vincent and M. Wayne Davis for their critical reviews of this manuscript. EMJ is an investigator at Howard Hughes Medical Institute.

## FUNDING

**Erik Jorgensen**

National Science Foundation NeuroNex 2014862 National Institute of Health NIH NINDS R01 NS034307

**Sean Merrill**

NIH F31 NRSA Predoctoral Fellows grant F31 NS084826

**Zhao-wen Wang**

National Institute of Health NIH R01 MH085927 and R01 NS109388

**Andres Villu Maricq**

National Institute of Health NIH R01 NS094421

## Author Contributions

BDM SAM EMJ Wrote manuscript

SAM SW BDM ZWW EMJ Designed experiments

SAM SW EMJ Conceived of the project

PMC AVM Performed behavioral recording and analysis

PL Performed electrophysiology

BDM PL Analyzed electrophysiology data

SAM BDM Performed single molecule localization microscopy

BDM LVD designed Proberuler

BDM SAM Analyzed single molecule localization data

SAM AC Performed genetic crosses

SAM AC BDM cloned plasmids

SAM BDM Generated transgenic animals

SW Performed and analyzed time resolved electron microscopy

BDM AS MS annotated serial reconstruction

AS MS Performed serial reconstruction electron microscopy

---

**Figure 2, Extended Data.**
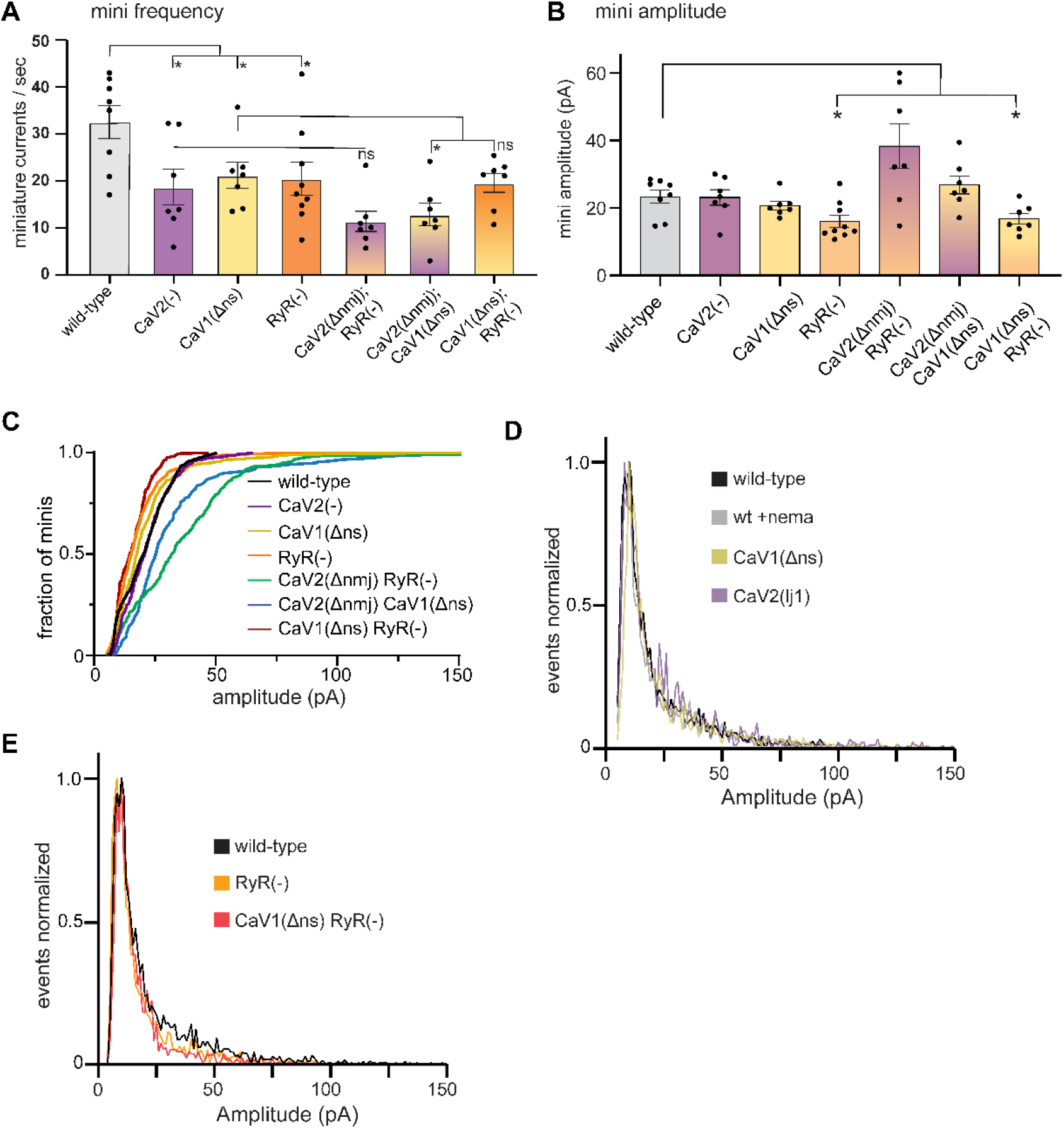
The ryanodine receptor acts in parallel to CaV2. (A) CaV2 and CaV1-RyR contribute additively to tonic release. Muscles were voltage-clamped, and tonic miniature synaptic currents recorded in 0.5 mM extracellular calcium: wild type: 33 ±4 minis/s, n=8; CaV2(−): 19 ±4 minis/s n=7, CaV1(Δns): 21 ± 3 minis/s n=7, RyR(−): 20 ±4 minis/s n=9. CaV2(Δnmj) RyR(−): 11 ± 2 minis/s n=7, CaV2(Δnmj) CaV1(Δns): 13 ±2 minis/s n=7, CaV1(Δns) RyR(−): 20 ±2 minis/s n=7. Welch’s t-test was used to calculate significance (B) RyR is required for large-amplitude spontaneous events. At 0.5mM calcium, wild-type 23 ±2 pA n=8, and CaV2(−): 23 ±2 pA n=7, CaV1(Δns): 21 ±1 pA n=7, RyR(−): 16 ±2 pA n=9. CaV2(Δnmj) RyR(−): (38 ±3 pA n=7, CaV2(Δnmj) CaV1(Δns): 26 ±3 pA n=7, and CaV1(Δns) RyR(−): 17 ±2 pA n=7. Welch’s t-test was used to calculate significance. (C) Cumulative distribution plot of mutant amplitudes. (D) Frequency distribution plot of voltage-gated calcium mutants amplitudes normalized to the mode. (E) Frequency distribution plot of ryanodine mutants with reduced amplitudes, normalized to mode. Error bars reported in SEM. *p<0.05, **p<0.005, ***p<0.001, ****p<0.0005 Data available as Source Data 2

**Figure 4, Extended Data.**
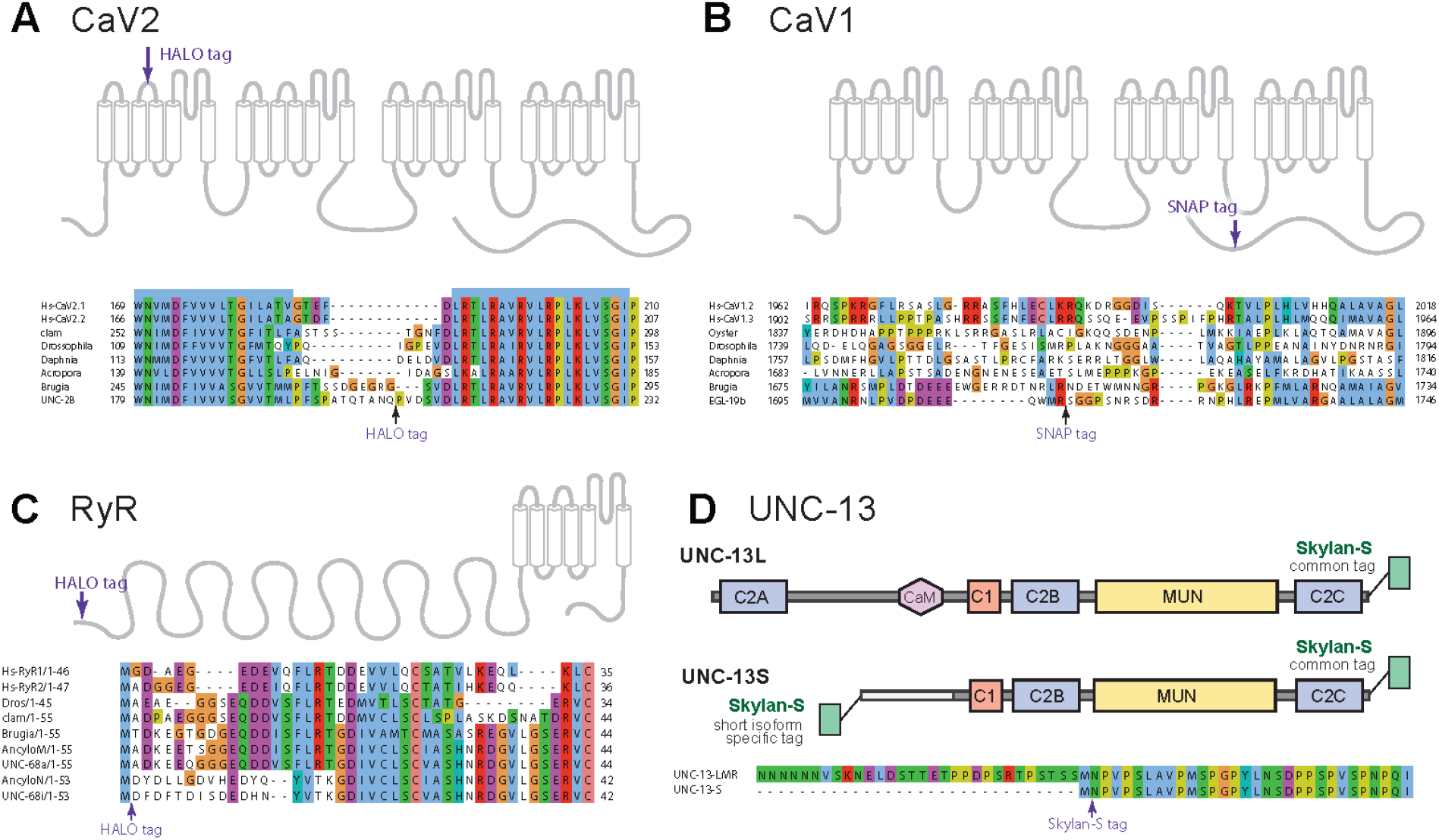
Tagging sites for calcium channels and UNC-13. (A-C), Tagging strategies and sites used for CRISPR/Cas9 tagging of the endogenous loci for CaV2, CaV1, and RyR. Regions with low conservation were targeted for insertion of tags into the genomic locus of each gene, *unc-2*, *egl-19*, and *unc-68*, respectively. (D) Tagging strategy at the endogenous locus of *unc-13* CRISPR / Cas9. The C-terminal tag labels all isoforms of UNC-13. The N-terminal tag labels the UNC-13S isoform. In addition, it will label a rare transcript UNC-13-LMR (∼2% of transcripts), that includes the C2A domain, sequences upstream of UNC-13S, and all sequences included in UNC-13S (wormbase.org version ws284).

**Figure 5, Extended Data.**
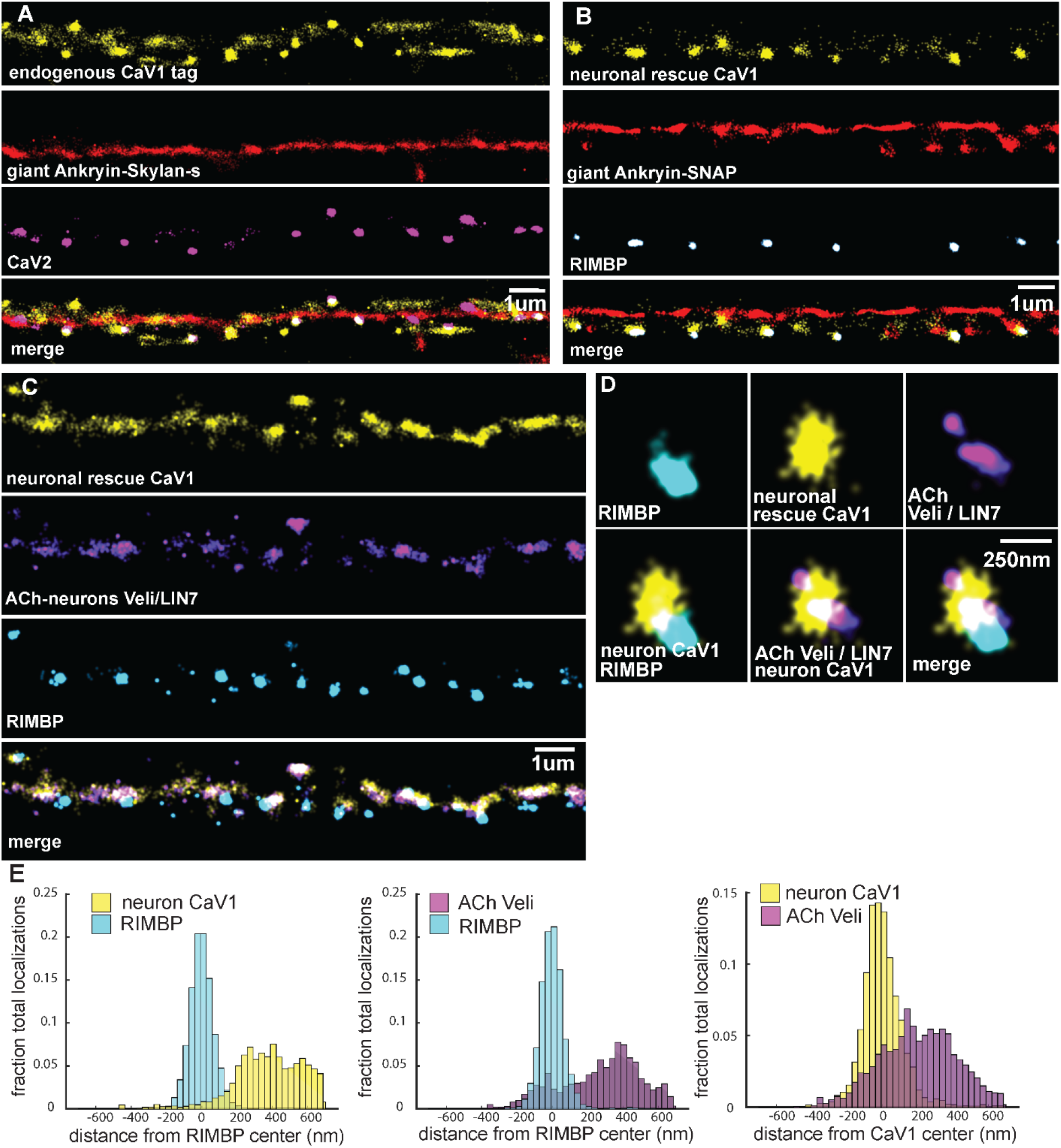
Tagged CaV1 expressed in neurons forms clusters at presynaptic boutons. Comparison of endogenously-tagged CaV1 and neuronally expressed rescue of CaV1. (A) Endogenous CaV1 tag: Localization microscopy images of dorsal nerve cord with CaV1-SNAP stained with STL-JF549cp (yellow), CaV2-HALO stained with HTL-JF646 (purple), and Giant Ankyrin-SkylanS (red). scale bar = 1um. (B) Exogenous CaV1 in the CaV1(Δns) strain. In the CaV1(Δns) background, CaV1 was rescued in neurons using a single copy transgene insertion of P*snt-1* promoter driving CaV1-HALO in neurons. Localization microscopy images of dorsal nerve cord. Dorsal cord of animals labelled with neuronal CaV1-HALO stained with HTL-JF646 (purple), Giant Ankyrin-SNAP stained with TMR-Star (red) and RIMBP-SkylanS (cyan). scale bar = 1um. (C) Neuronal CaV1 colocalizes with Veli/LIN7 expressed in acetylcholine neurons. In the CaV1(Δns) background, CaV1 was rescued in neurons using a single copy transgene insertion of P*snt-1* promoter driving CaV1-HALO in neurons and stained with HTL-JF646. Veli / LIN7 was tagged with SNAP and stained with STL-JF549cp, and was expressed in acetylcholine neurons using the *unc-129* promotor as an extrachromosomal array. scale bar = 1um. Localization microscopy images of dorsal nerve cord. Dense projections are marked by RIMBP-SkylanS. (D) Single synapse analysis of Veli / LIN7, CaV1, RIMBP. Dense projections are marked by RIMBP-SkylanS. In the CaV1(Δns) background, CaV1 was rescued in neurons using a single copy transgene insertion of the P*snt-1* promoter driving CaV1-HALO, and stained with HTL-JF646. Veli/ LIN7 was expressed in acetylcholine neurons using the P*unc-129* promotor as an extrachromosomal array, and tagged with SNAP and stained with STL-JF549cp and. Scale bar = 250nm. (E-G) Cumulative distribution plot of P*snt-1*:CaV1-HALO to RIMBP-SkylanS center, and Veli / LIN7 tagged with SNAP to RIMBP-SkylanS center measured from synaptic regions. n=24 synapses, N=5 animals. Data available as Source Data 5

